# Replay as structural inference in the hippocampal-entorhinal system

**DOI:** 10.1101/2020.08.07.241547

**Authors:** Talfan Evans, Neil Burgess

## Abstract

Model-based decision making relies on the construction of an accurate representation of the underlying state-space, and localization of one’s current state within it. One way to localize is to recognize the state with which incoming sensory observations have been previously associated. Another is to update a previous state estimate given a known transition. In practice, both strategies are subject to uncertainty and must be balanced with respect to their relative confidences; robust learning requires aligning the predictions of both models over historic observations. Here, we propose a dual-systems account of the hippocampal-entorhinal system, where sensory prediction errors between these models during *online* exploration of state space initiate *offline* probabilistic inference. *Offline* inference computes a *metric* embedding on grid cells of an *associative* place graph encoded in the recurrent connections between place cells, achieved by message passing between cells representing non-local states. We provide testable explanations for coordinated place and grid cell ‘replay’ as efficient message passing, and for distortions, partial rescaling and direction-dependent offsets in grid patterns as the confidence weighted balancing of model priors, and distortions to grid patterns as reflecting inhomogeneous sensory inputs across states.

**Author Summary:**

- Minimising prediction errors between transition and sensory input (observation) models predicts partial rescaling and direction-dependent offsets in grid cell firing patterns.
- Inhomogeneous sensory inputs predict distortions of grid firing patterns during *online* localisation, and local changes of grid scale during *offline* inference.
- Principled information propagation during *offline* inference predicts coordinated place and grid cell ‘replay’, where sequences propagate between structurally related features.

## Introduction

Grid cells in the medial entorhinal cortex (mEC), whose firing fields form a periodic hexagonal lattice across the environment, are thought to support path integration^1–3^, whereas hippocampal place cells tend to have unimodal firing fields reflecting environmental cues such as boundaries^4,5^. Grid cell firing patterns are stable over time, suggesting corrective environmental inputs, possibly from place cells^6^, but rely more on self-motion than place cell firing patterns^7^ suggesting that environmental inputs are not fully corrective^2,8,9^. Given estimates of each input’s uncertainty, Bayesian inference tells us how they should be optimally combined.

Although *online* learning (i.e. using only currently available sensory information) can converge under low PI and sensory noise, robust learning in the presence of noise requires minimizing the error between self-motion and environmental estimates of location across all state transitions^10,11^. Thus, historic observations must be stored and revisited *offline* (i.e. independently of current sensory inputs) to allow propagation of local environmental information to non-local but structurally connected regions of the cognitive map, e.g. as when adapting to a novel shortcut or barrier. This process can also be viewed as an embedding of sensory experience within a low-dimensional manifold (in this case, 2D space), as observed of place cells during sleep^12^.

Building on previous work^11,13–15^, we propose a dual-systems *(online-offline)* account of spatial inference in the hippocampal/entorhinal system, which we define as the process of identifying the configuration of both one’s own location (current state) and the location of environmental landmarks in space (c.f. ‘SLAM’^10^). In familiar environments, *online* localization (identification of one’s own position) is achieved by recursively combining self-motion and sensory inputs, which are mediated by learned *transition* and *observation* models, respectively. However, prediction errors between these models trigger *offline* inference over non-local states, facilitating fast learning of new or changed associative environmental structure, encoded *online* in place-place cell synaptic associations. We identify this *offline* inference with coordinated hippocampal/mEC ‘replay’^16–20^.

Our framework also provides algorithmic- and implementation-level explanations for observed features of grid cell firing in response to manipulations^7,9,21–24^ or inhomogeneity^25–29^ of environmental sensory input. Overall, these phenomena can be understood in a probabilistic framework, where minimization of prediction errors between the transition and observation models are traded against prior model beliefs.

## Results

### Probabilistic *online* localization with place and grid cells

Grid cells exist in ‘modules’ of cells, whose firing patterns have the same spatial scale and orientation relative to the environment, but differ in their spatial offsets^30^. The spatial scale increases in discrete steps along the dorso-ventral axis, suggesting that, across modules, GCs support a hierarchical representation of space^21,24,31–33^. Here, we consider a single module of GCs, whose activity represent a probability distribution over a periodic, discretized region of space (visualised as a topographically arranged sheet of cells; Fig. 1A).

**Figure 1.**
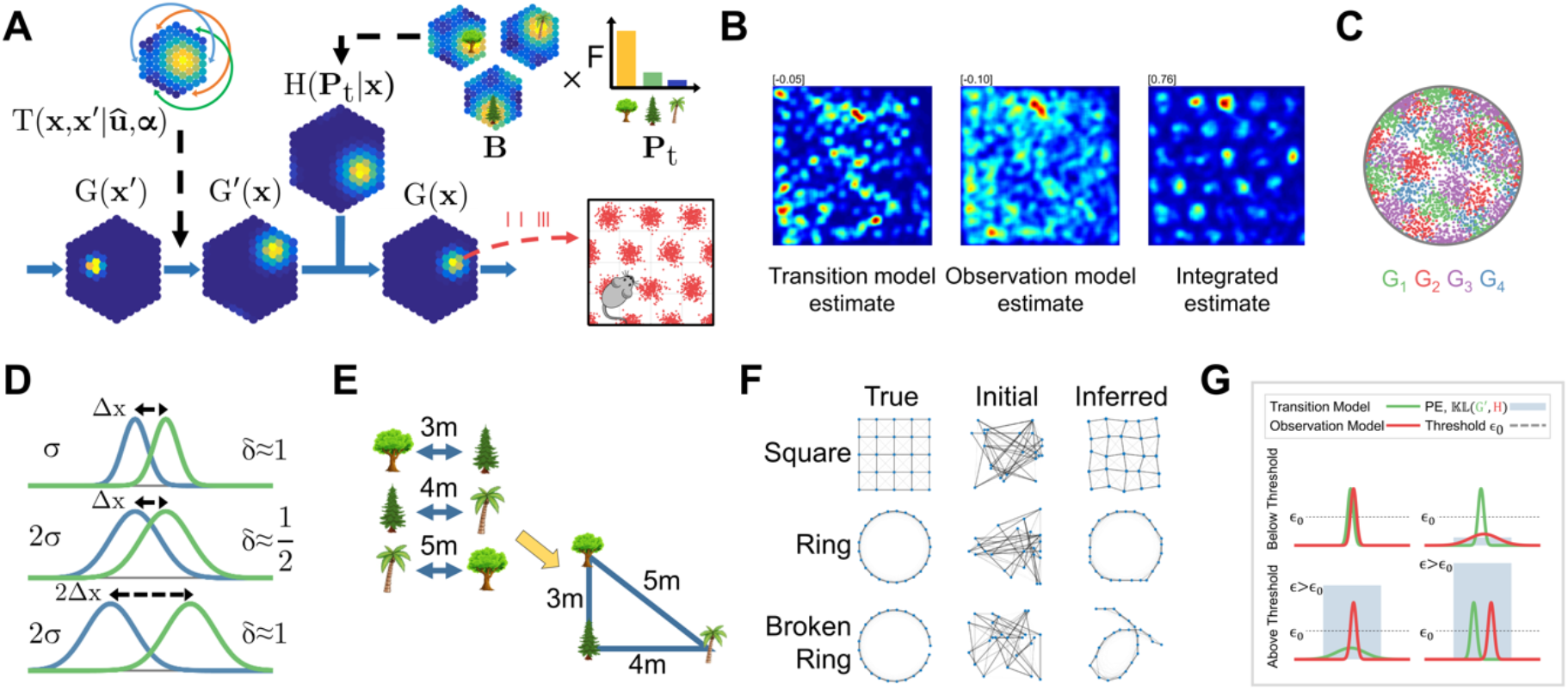
Online+offline localization and mapping. **A)** Illustration of recursive Bayesian integration. A probability distribution over current location (*G*(***x***)), represented by grid cell firing, is updated according to self-motion via the *transition model* (T(**x,x**’|**û, α**)) then refined by environmental inputs via the *observation model* (*H*(***P_t_*** |***x_t_***)). **C)** Estimates based on the integration of noisy self-motion and environmental inputs may be stable, as shown in simulated grid cell firing rate maps *(right),* despite instability when using only self-motion *(left)* or environmental inputs after brief initial exploration *(middle;* numbers show gridness score). **C)** Simulated grid cells exhibit spatially offset grid-like firing patterns, due to toroidal connectivity, despite the absence of attractor dynamics. Right shows histogram of spatial phases. **D)** Inferred pairwise distances D are a function of the ‘overlap’ between place fields. **E)** Pairwise distances can be used to infer the structure of the world (the mapping of place fields onto the grid map). **F)** Given noisy initial priors (“Initial”), structural encodings are modified to reflect pairwise *associative* measurements. Inferred structure is sensitive to the topology of the environment (cf. “Ring” and “Broken Ring”). **G)** Illustration of the prediction error mechanism used to arbitrate between the *online* and *offline* systems (blue bars show prediction error ε, ε_0_ is the minimum prediction error needed to trigger *offline* inference).

The self-location distribution is maintained over time by recursively integrating sensory and selfmotion inputs, accounting for their uncertainties (Fig. 1A). Firstly, the posterior distribution over agent location (grid module activity) from the previous time-step ***G*** is updated given noisy perceived movement ***û*** via the *transition model T* (see Methods):

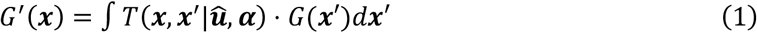

where ***x*** is the 2D coordinate of the agent location in *metric* space (corresponding to a particular grid cell) and ***x***’ the location at the previous time-step. Biophysically, *T* would be represented by a population of direction dependent ‘shifter’ cells with asymmetric recurrent weights^1^ with a circulant structure^34^ (Fig. S1C, see Methods) learned *apriori* (but see Refs. ^35–37^). The rate of translation of activity on the grid sheet in response to movement ***û*** is controlled by the *transition model* gain ***α*** = [*α_x_*, 0; 0, *α_u_*], which might correspond to the strength of the associations to, or the speed dependence of, shifter or conjunctive cells^31,38,39^ (see Supplementary Methods).

The *transition model* estimate *G*’(***x***) is then refined by observations of environmental features, which map to *metric* space locations via *observation model H:*

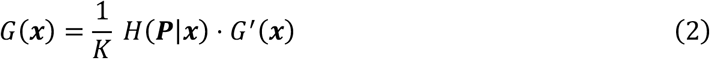

where *P* is a vector of place cell firing rates, firing of place cell *i* representing the likelihood of the presence of a specific sensory feature, or combination thereof. In our simulations, these have unique locations in physical space **μ_i_** and receptive field widths 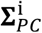. Where the number of grid cells is large, the weights from place cell *i* to the grid module define a distribution for that feature’s estimated location in *metric* space (Fig. 1A). The weighted projection of place cell activity by these weights defines the *observation model H. K* provides inhibitory normalization (see Supplementary Methods).

*Online* learning modifies the *observation model* to reflect the current *transition model* location estimate (induces synaptic changes in the place-grid cell connection weights via a BCM rule; Fig. 2A; Methods). *Online* learning produces stable grid patterns (due to the circulant structure of *T)* for a range of levels of PI and sensory noise, but convergence fails in higher noise regimes (Fig. S2B). After a short period of initial learning, stable grid patterns emerge in the integrated estimate, despite the pure PI estimate being too noisy and the sensory associations too immature to drive stable patterns, if operating independently (Fig. 1B).

**Figure 2.**
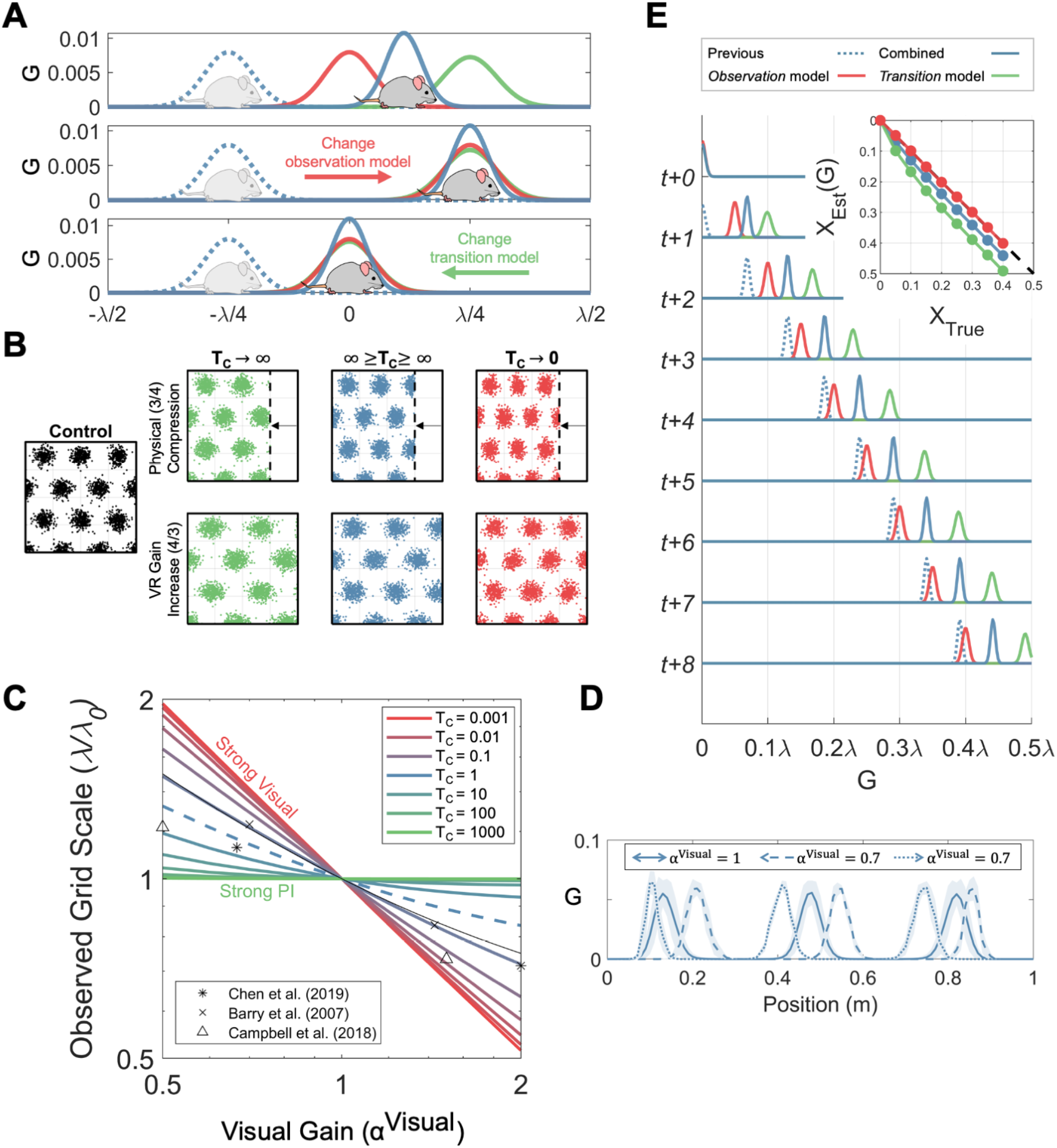
Minimizing prediction errors in the *offline* system: grid rescaling and direction-dependent offsets under manipulations of environmental size or VR gain. **A)** During *online* spatial localization, the *observation* model estimates location in metric space (activity on the grid sheet) via inputs from place cells driven by environmental features (red curve), the *transition* model updates the previous estimate (dashed blue curve) according to self-motion (green curve), producing a combined estimate (blue curve). Manipulations of the environment cause the predictions from both models to diverge. One way to minimize these prediction errors is to modify the *observation* model by changing the connection weights from place to grid cells (i.e. the mapping between environmental observations to estimated location in grid space). An alternative is to modify the *transition* model to reflect the *observation* model estimate (e.g. varying the ‘gain’ mapping self-motion to grid space). The degree to which both are modified is controlled by the relative strength of their respective prior confidences (T_c_). **B)** Joint optimization of the *observation* and *transition models* predict partial rescaling of grid patterns in response to increase in the VR gain (i.e. the rate of visual movement in response to physical movement on the ball; below) or compression of a real environment (above). When the system is confident in its self-motion (T_c_ → ∞, green), the *observation* model is modified to match the *transition* model (no change in grid scale plotted in the real or visual VR environment). When the system is confident in its environmental inputs (T_c_ *→* 0, red), the *transition* model adapts and grid scale follows the environmental change. Balanced model confidence produces intermediate rescaling (blue). **C)** Change in observed grid scale (grid pattern plotted against self-motion) depends on the *transition* confidence values T_c_. X-axis shows VR gain change or environmental compression, where α^visual^ = 1 is a control trial. Y-axis shows observed change in grid scale (see Methods). Data points show corresponding values from Barry et al. (2007), Chen et al. (2019) and Campbell et al., (2019), which suggest an approximately equal weighting of *transition* and *observation* model priors for both gain decreases and increase trials (T_c_ ≈ 1). **D)** Firing rate map of a grid cell on the linear track in control (solid line) and VR gain decrease trials (*α^visual^* = 0.7), plotted in the visual VR environment. In the gain decrease condition, the grid fields are stably shifted to fire earlier when running left (dashed line) or right (dotted line), i.e. towards the location indicated by the transition model from that indicated by vision. **E)** The directiondependent shift in estimated location in the grid module in a VR gain decrease trial (*_α_^visual^* = 2/3) stabilises at a fixed distance. Sequence of eight updates of estimated location on the grid module G(x) when running to the right in a VR visual gain decrease trial, colours as in A. The *transition* model (green) predicts a location ahead of that from the *observation* model on the grid sheet (red; driven by visual input) because of the visual gain decrease. Combining these estimates produces an intermediate distribution (blue). At each new update, the prediction from the *transition* model builds on the shift of the previous combined estimate (not the previous transition model estimate) so that the distance between the observation model estimate and the combined estimate stabilizes at a fixed value, producing a fixed direction-dependent offset of the grid patterns in environmental coordinates (see the differences in the location estimates between models, Inset).

### *Offline* inference: The hippocampus as a probabilistic graph

Local, *online* learning is not robust in novel environments, because corrections to the estimated agent location (current grid cell activity), e.g. upon encountering familiar environmental features (place cell activity) associated to a different location on the grid module, also imply corrections to the encoding of feature observations along the preceding trajectory^10^. That is to say; local updates to the *cognitive map* also imply non-local, structurally associated changes. Formally, probabilistic spatial inference in this case requires finding the most likely configuration of *metric* space feature locations {**b**_*i*_}_*i*=1:*N_p_*_ (each ***b_i_*** is the 2D coordinate of feature *i* in *metric* space, i.e. a place - grid cell association) and agent location *x* (the distribution over which is indicated by the grid cell firing rates) consistent with environmental sensory observations made along a given trajectory.

Theoretically, the configuration {***b_i_***}_*i*=1*N_p_*_ can be recovered purely from the distances between pairs of environmental features^10^ (Fig. 1F, *“square”, “ring”).* Importantly, despite both feature locations being susceptible to large absolute errors (due to noisy PI), the errors will be correlated such that pairwise distance measurements will decrease in variance with observations^10^. This method predicts characteristic failure modes when the pairwise distance information is ambiguous or incomplete (e.g., Fig. 1F; *“broken ring”).* New distance observations might also cause dramatic changes to the inferred configuration (e.g. the discovery of a shortcut). If the current *absolute* estimates of feature node (i.e. place cell) location are stored in the place-grid cell synaptic weights, we propose that the *relative distances* between pairs of features are stored in the recurrent weights between place cells in hippocampal region CA3.

Consider a spring network, where the edge between environmental feature nodes *i* and *j* represents a noisy pairwise observation with length reflecting pairwise distance and stiffness reflecting certainty^13,40^ (Fig. 1F, S6A). Minimizing the elastic energy in the spring mesh system corresponds to finding the maximum of the joint likelihood *L*(·), which is a function of the feature locations in *metric* space {***b_i_***}_*i*=1:*N_p_*_ and the internal gain parameter ***α***, given pairwise distance measurements with Gaussian noise (*δ_ij_*). This is equivalent to defining a probabilistic graphical model (see Methods) over the posterior:

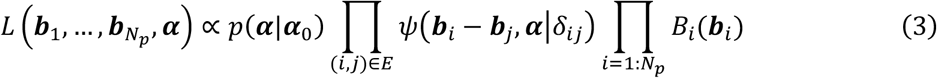

where the current PC-GC weights *B* (*B_i_* is the probability distribution of the location of feature *i* and *B_i_* (***b**_i_*) is its value at *metric/grid* module location ***b**_i_*) act as priors on the feature node locations, the pairwise potential terms *ψ*(·) penalize the difference between *associative* pairwise distance measurements *δ_ij_*, made directly in environmental *stimulus* space, and the distance between their candidate locations in *metric* space ***b_i_ — b_j_***. *E* is the set of connected PCs (see Methods). Distance in *metric* space is also a function of the *transition model* gain (***α***), which has a Gaussian prior *p*(***α|α***_0_) (a larger |***α***| will decrease the *metric* space distance for all pairs; see Methods). Maximizing the likelihood (finding the state of minimum energy in the spring network) model over all feature node pairs minimizes the total prediction error between *associative* and *metric* generative models of the world^15^ (Fig. S7).

The *associative* distances can be straightforwardly learned during *online* exploration. Since Hebbian learning reflects coactivity, a trajectory exploring the environment uniformly results in synaptic strengths between place cells proportional to the spatial correlation between their receptive fields^41^. The Euclidean separation between their fields is then accessed via a simple transformation (see Methods; Fig. S1G). In this context, learning the PC-GC weights (modifying the *observation* model) during *online* localization corresponds to forming spatial priors over feature locations which anchor the structure, which would otherwise be translation or rotation invariant (since measurements are relative), learned during *offline* inference to constant locations on the grid-map. Taken together, our framework proposes a mapping onto anatomy of the joint *agent-feature* location distribution required for full probabilistic inference over environmental structure (Fig. S5; See overall algorithm in Table S1).

### Partial grid pattern responses to environmental rescaling

Uniform rescaling of an environment will introduce a mismatch between the estimates of location from the *transition* and *observation* models (i.e. a ‘prediction error’). To minimize these prediction errors, the *offline* system can either modify the *transition* model gain to match the current environmental input (Fig. 2A, bottom), or modify the mapping from environmental inputs to *metric* space in the *observation* model (Fig. 2A, middle; see Methods). The degree to which either is modified should reflect their relative confidences, specified by a ‘transition confidence score’ (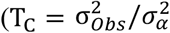, the ratio of confidence in the *transition* vs *observation* models; see Methods). Similarly, a ‘prior confidence score’ P_c_ specifies how much the system will tolerate persistent prediction errors; if P_c_ is large, optimization may favour preserving prior configurations, as opposed to alignment of the current *transition* and *observation* models (see Methods).

We modelled experiments in which the physical environment^21,42^ and perceived velocity through a visual virtual environment^7^ were re-scaled, such that self-motion and sensory inputs conflicted. In both experiments, the rescaling of the grid patterns was *partial,* i.e. less than the magnitude of the physical or virtual manipulation, and less than those of place fields.

Both manipulations can be simulated by introducing a visual gain parameter *α^Visual^* to the simulation of the environment (in both experiments it scales the amount of self-motion required to traverse the width of the perceived environment). Learned *associative* distances (***δ***) are also scaled by this parameter, reflecting its effect on the temporal overlap of place fields (see Methods Eq. 12). We simulated grid pattern rescaling responses over a range of transition confidence scores T_c_. When confidence in the *transition* model is high (T_c_ → ∞), grid patterns in the real world are unchanged when plotted against physical movement (but are changed when plotted in visual VR coordinates; Fig. 2B, first column). The opposite is true when confidence in the *observation* model is high: grid patterns are unchanged relative to the apparent environment (Fig. 2B, last column).

However, for intermediate T_c_ values (i.e. balanced confidence in the *transition* and *observation* models), the model predicts partial rescaling of the grid pattern relative to the size of the manipulation (Fig. 2B, middle column), matching the observed grid patterns in both experiments^7,21^ (and a similar third experiment in on a virtual linear track^43^; Fig. 2C).

### Differential grid and place field responses to environmental reshaping

How does the *offline* system respond to more complex environmental deformations? When one wall of a familiar rectangular environment is rotated inwards by 45°^22^, place fields near the wall shifted almost fully while fields further away remained largely stationary, consistent with place fields preferentially reflecting local environmental inputs^5^. In contrast, grid fields shifted only partially near to the manipulated wall. Using the observed place field shifts^23^, we simulated the response of grid cells to the same manipulation (Fig. 3; see Supplementary Methods). Shifted place fields induce a misalignment between the *associative* distances and the distance between their encodings in *metric* space. The place field shifts are local and non-uniform, and so misalignment cannot be corrected by a global change to the *transition gain **α**.* Indeed, ***α*** is not significantly modified during the optimization process, regardless of the T_C_ value. Instead, alignment between the *transition* and *observation* models is maximized by modifying the *observation model,* i.e. updating the locations of the place fields on the grid module.

**Figure 3.**
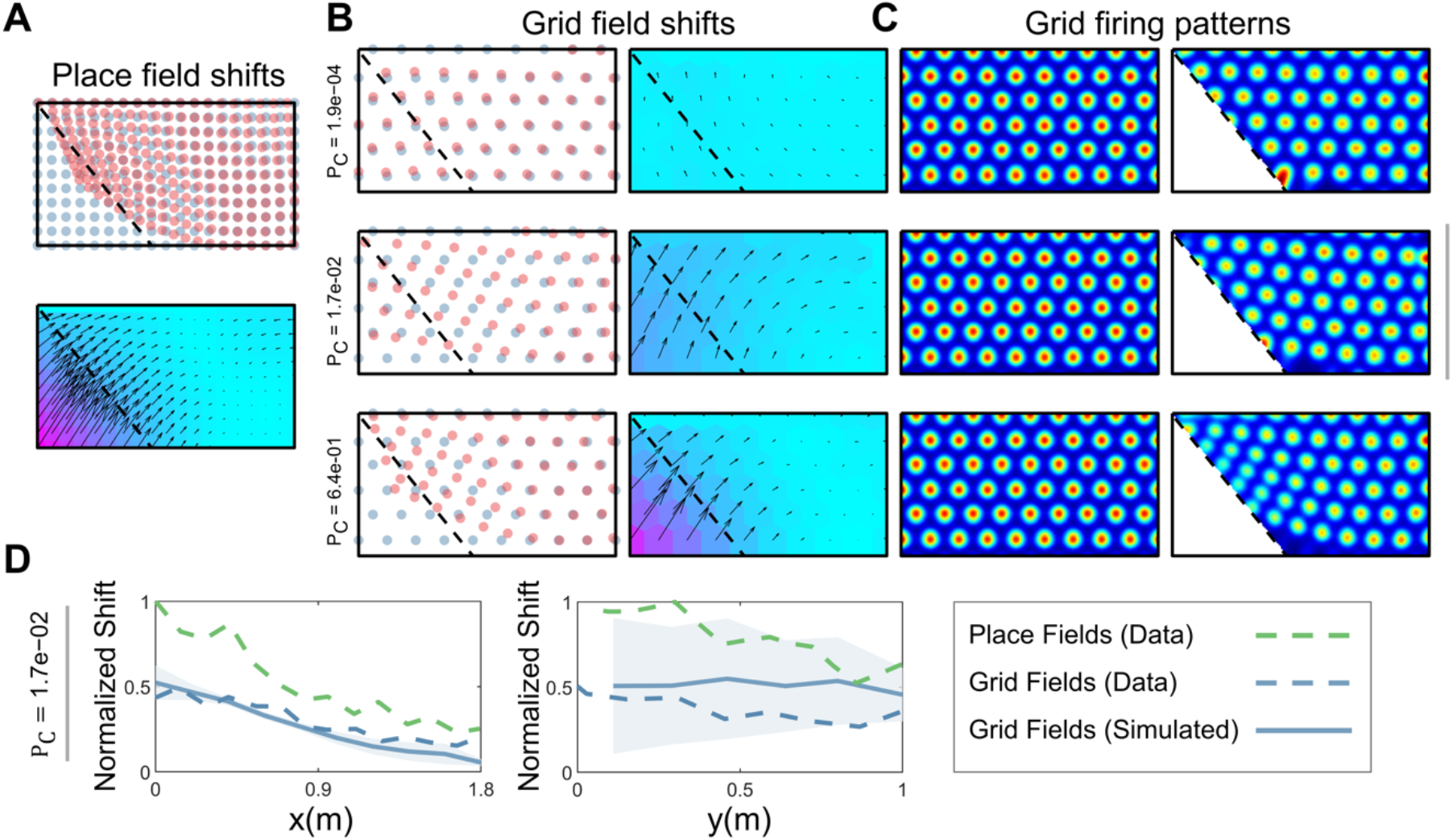
Effects of local deformation of a rectangular environment on grid patterns following *offline* inference (Krupic et al., 2018). Dots indicate place/grid field locations, arrows indicate shifts before/after *offline* structural optimization and coloured hexagons indicate magnitude of shift vector. Grid patterns show estimate generated by the *observation* model (weighted place cell activity). **A)** Place field shifts were measured from Krupic et al. (2018) and interpolated and smoothed. **B-C)** Place field shifts cause immediate prediction errors (PE) between the new pairwise place field distances and the distances between their corresponding grid locations. If the *observation / transition* model priors are weak (P_c_ → 0), PEs are eliminated during *offline* inference by updating the *observation* model (top row; unlike in Fig. 4B, modifying the global gain did not remove the effect of local distortions). Complete adaptation in the *observation* model leads to an unperturbed grid pattern, i.e. grid fields will not shift. Alternatively, strong model priors (P_c_ *→ ∞)* prevent adaptation to new environmental inputs, leading to distorted grid patterns when driven purely by the observation model, whose field shifts match those of the place cells (bottom row). When the model priors are balanced against the new pairwise observations, the *observation* model is partially adjusted, producing partial grid field shifts (which are smaller than those of the place fields; middle row). Partial (middle) or no (bottom) adjustment to new observations preserve mismatches between the *transition* and *observation* models, which would result in direction-dependent offsets (see Figure 4E) and irregular firing patterns (see Figure S2D). NB the confidence in the prior model may depend on location, e.g. if there is strong anchoring to the wall prior connections from place cells with fields near the wall may be stronger than those with fields further away. **D)** Comparison of experimentally observed place and grid field shifts with simulation corresponding to the middle row of **B-C**.

If there were no confidence in the prior *observation* model (P_C_ → 0), it would be modified offline to match the transition model, leaving the grid pattern unperturbed by the environmental change (Fig. 3B,C, top row). At the other extreme, favouring prior beliefs over recent observations (i.e. pairwise distances encoded during the manipulation trial) would result in an unchanged *observation* model, and grid field shifts that exactly mirror corresponding place field shifts (Fig. 3B,C, bottom row). In this regime, there would be permanent misalignment of the *transition* and *observation* models during *online* localization, producing noisy grid patterns, as when simulating a related experiment where grid distortions were observed in trapezoid environments^44^ (Fig. S2D). Setting P_C_ to an intermediate value reproduces the experimentally observed partial shifting of grid fields (relative to the place fields^23^) when visualizing the structure encoded in the *observation* model (i.e. assuming low confidence in the transition model; Fig. 3B,C and D).

### Direction dependent shifting of grid patterns during *online* localization

In addition to partial changes to grid scale in response to environmental rescaling, enduring misalignments between *observation* and *transition* models can result from strong model priors, which prevent complete adaptation of the *transition* model gain. These cause the *transition* estimate to consistently precede that of the *observation* model, in the current direction of travel (on a 1D track, Fig. 2E) during VR visual gain decrease trials or physical expansion of the environment.

In all three cases, the integrated estimate of location (Eq. 2) in the *online* model converges to a fixed distance ahead of the *observation* model estimate (in the direction of travel; Fig. 2E inset and 3D), causing the grid pattern in the real world to dynamically shift opposite the direction of travel, as observed experimentally^7,9^. Our model suggests that the offsets should be partial (smaller than implied by a hard-reset at the boundary) and not specifically require a recent boundary encounter (cf. Keinath et al.^9^). Dynamic shifting in the model will reduce with experience of the novel or manipulated environment, as model misalignment reduces, as observed experimentally^45^.

### *Online* and *offline* perceptual warping in spatial representations

With increasing experience of an environment, grid firing patterns exhibit both local scale changes^26^ and global shear-like distortions^29^, the latter associated with 7.5-8° offsets of one of the grid axes^29,44^ to the walls of square environments. Both effects were present in our simulations and can be attributed distinctly to the *offline* map-learning and *online* localization components of our theoretical framework.

Firstly, we show that local changes to the grid scale^26^, which are positively correlated with behavioural occupancy (animals spend more time in the middle of the environment), arise from the *offline* process of map (PC-GC connections) learning. These mapping-induced distortions can be further subdivided into two mechanisms, both of which induce local scale changes by biasing the pairwise distances recovered from the Hebbian learned recurrent connections in CA3 (Fig. 1D,4; Fig. S7).

**Figure 4.**
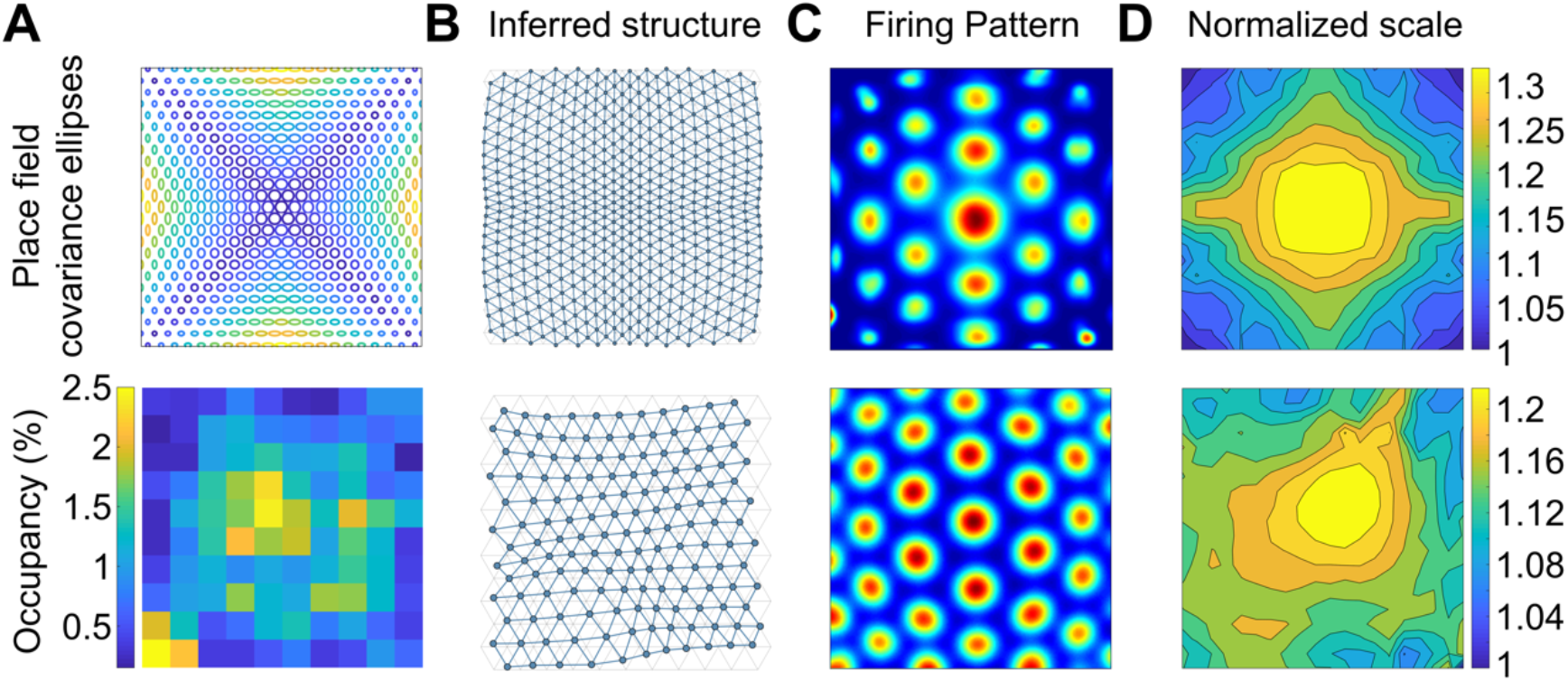
Distortions to grid patterns caused by inhomogeneous environmental input during offline inference. **A)** Simulated variation in place field shape due to proximity to boundaries (above), or inhomogeneous sampling of locations (Hagglund et al,. 2019; below) results in distortions to the inferred pairwise distances in CA3 (**B**). These distortions lead to distortions to grid scale due to adjustment of the observation model during offline inference (**C-D**).

Firstly, relative behavioural under-sampling of the place fields near the boundaries of the environment (using occupancy statistics from^26^; Fig. 4A, bottom) lead to weaker PC-PC connections, and consequent overestimation of their pairwise distances, producing local scale changes (Fig. S7C).

Secondly, since Hebbian learned connection weights between place cells reflect the correlation in their firing, and therefore their statistical discriminability^46^, two place cells with broad receptive fields would develop a stronger connection than a pair with equal separation but narrower receptive fields (stronger connections correspond to shorter distances on the grid module, producing grid patterns with larger scales in the environment; Fig. 1D). Another recent study^47^ suggests that place fields are narrower near the edges of an environment, consistent with greater precision when driven by more proximal environmental features^5^ (Fig. 4A, top row). In our model, this produces weaker recurrent connections and a shrinking of the grid pattern at the edges of the environment following *offline* inference (Fig. 4D,E, top row).

Together, our results suggest that the cognitive ‘distance’ between two sensory features should be greater both when the absolute confidence in their spatial locations is greater (reflecting an increased statistical discriminability), or when those features are under-sampled relative to other features.

Although the action of both mechanisms are independent their effect is the same; both i) relative under-sampling of the transition between two adjacent states and ii) a reduced statistical discriminability between those states, both contribute to a weaker pairing of their representative place cells, resulting in greater separation between their encodings in *metric* space and a locally larger grid scale when ‘read-out’ in the firing pattern (a locally larger perception of distance).

In contrast, global shear-like distortions^29^ and associated 7.5-8° offsets of one of the grid axes^29,44^ can be interpreted as *localization* induced distortions during *online* exploration. In Stensola et al.^29^, rats were introduced into the same corner of the box at the start of each trial; in Butler et al.^25^, shearing developed following the introduction of reward^25^. In both experiments, shearing developed with increasing experience^25,29^. We hypothesized that these distortions reflect an increasing effect of non-uniform environmental inputs to the grid module, either reflecting their natural distribution^25,29^ or inhomogeneous behavioural sampling of environmental locations^26^.

In our simulations, given a learned map, biasing the strength of sensory inputs at specific locations (e.g. one/two corners) during *online* exploration reproduced several experimentally characterized global distortions by causing a bias in the decoding of location (i.e. salient locations contribute a larger ‘vote’; Fig. S3; see Supplementary Methods).

### Probabilistic inference through HPC-mEC message passing

To this point we discussed, from a functional perspective, how the brain might optimize its internal representations to reflect the uncertainty of sensory information. But how might the brain perform this optimization? In the above analyses of *offline* inference, we numerically computed the maximally likely feature locations on the grid module. However, the system must also track the uncertainty in these estimates, which would require updating the place-grid cell weights (including those with firing fields far from the agent location). An update of the full weight distributions is generally intractable when the state space is large.

Belief propagation^48^ is a technique for approximating this inference on graph structured data, and comprises two stages. First, a given feature node (i.e. a place cell) computes its location distribution (i.e. connections to the grid cells) *B_i_* (***b**_i_*) by multiplying its prior 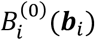 with messages received from its connected neighbours (Fig. 5C; see Methods for details). A message *m_i→j_* (***b**_j_*) expresses neighbour node *i*’s belief of node *j*’s location, conditioned on its own distribution, and is dependent on the same pairwise potential terms *ψ*(·) in Eq. 3. The effect of a message is to favour distributions of nodes *i* and *j* which locate them at a radial distance equal to the associative distance *δ_ij_*; causing messages to be expressed as rings centred on the belief of the broadcasting node (Fig. 5C). Resolving a feature’s unique location then depends on aggregating messages from multiple neighbours (Fig. 5C). Computations are distributed, and importantly only require information that is local to each neuron.

**Figure 5.**
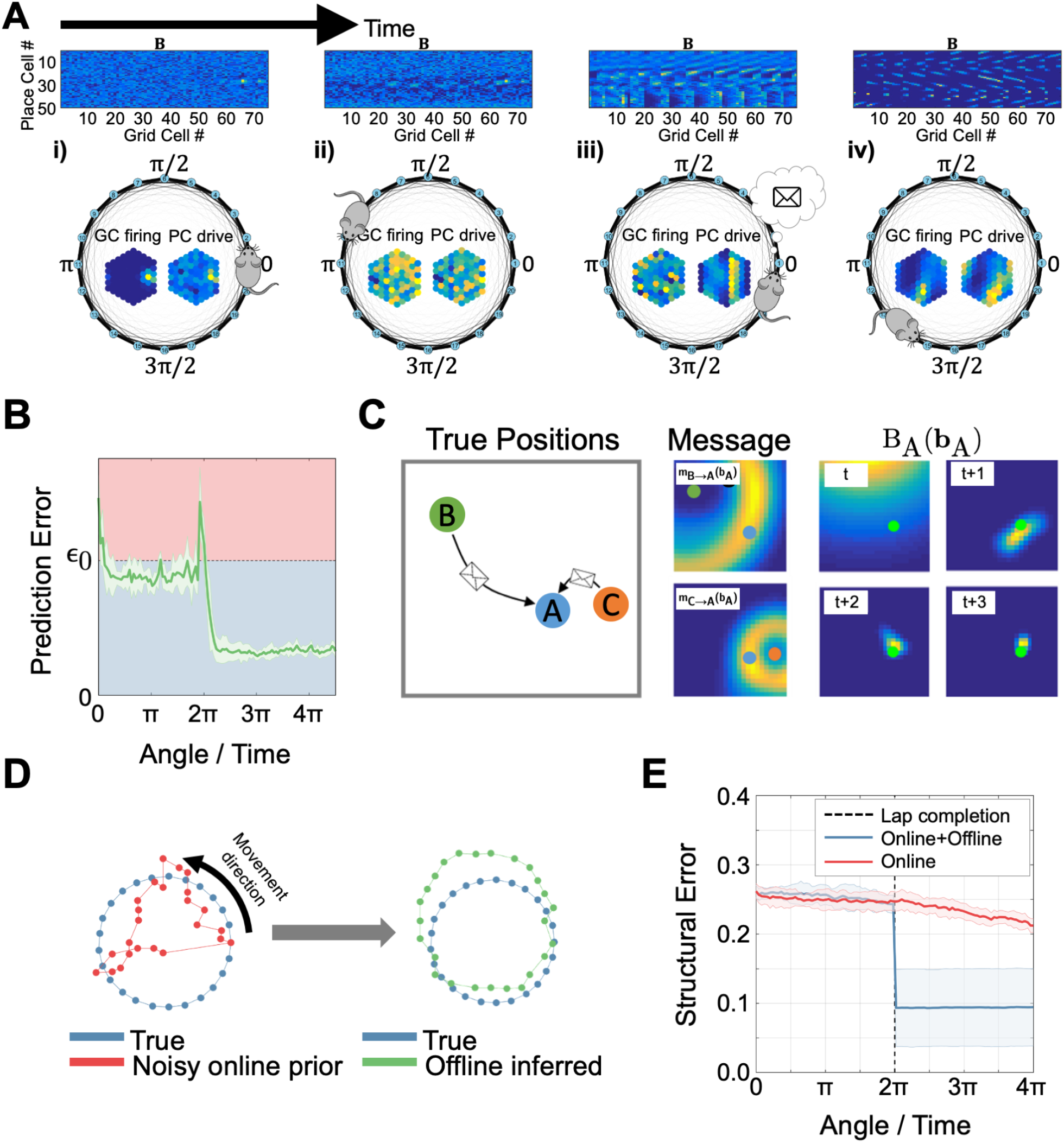
Illustration of the dual-systems model: prediction errors and replay in the loopclosure task (see also Supplementary Video 1). **A)** Place cell – grid cell (PC-GC) connection weights (above) as agent runs around a circular track for the first time (below, GC activity and the input from PCs to GCs, both shown on topographically organised sheet of cells, inset). **i)** Confidence in the initial location is high, such that coactive GC and PC fields form strong associations (the GC firing distribution is peaked and the inputs from place cells with fields at the beginning of the track are strong). **ii)** The agent navigates around the track, accumulating self-motion error, leading to diffuse GC firing. **iii)** Prediction errors (PE) on lap completion (when the initially learned precise PC input arrives) triggers an *offline* inference event (see main text and Supp. Video 1 for details). **iv)** On subsequent laps of the track, PC-GC weights are sharply tuned following *offline* inference, allowing effective localization. **B)** PE is reduced on completion of subsequent laps due to alignment of the *transition* and *observation models* (i.e. environmental inputs and self-motion updating of GC activity coincide). **C)** Illustration of belief propagation. Place cell A receives messages from PCs B and C. Messages take the form of rings, describing a preferred distance about the current locations of B and C with variance reflecting the confidence in the message (the variance of pairwise distance estimates with Gaussian noise). The intersection of the messages uniquely determines the location of A over time. NB A will also be broadcasting messages back to B and C. **D)** True structure (blue), structure encoded by noisy path integration (left, red; i.e. the location of the peaks of the weights from each place cell to the grid cell sheet) and structure inferred after loop-closure (right, green). **E)** *Offline* inference allows one-shot learning when compared to the online system alone.

Each node in the graph iterates between updating its belief and broadcasting messages, converging when new messages cease to change the beliefs of their recipient nodes. As expected, the reduction in pairwise prediction error between associative distances and their corresponding distances in grid space (see Methods) over successive message iterations is accompanied by a sharpening of the distribution of each feature’s location on the grid module (see Supplementary Methods; Fig. S1H).

### *Offline* inference triggered by prediction errors

How might the *online* and *offline* systems interact? If the *online* system is sufficient to localize within pre-learned, simple or slowly changing environments, non-local reactivations of place cells would be unnecessary. However, more complex *offline* inference is required under more demanding circumstances, or in novel or changing environments. We hypothesize that *offline* or ‘remote’ inference is triggered by *prediction errors* between location estimates from the *transition* and *observation* models, respectively (Fig. 1G), defined in our model as the Kullback-Leibler divergence 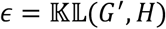.

Prediction errors are large when the *observation model* prediction (weighted place cell input) is different and more sharply peaked than the *transition model* estimate (Fig. 1G; see Methods; prediction errors will not be generated in absence of incoming sensory information, as in darkness, when the observation model estimate is uncertain).

To illustrate our *dual-systems (online+offline)* hypothesis (Fig. S7), we simulated an agent navigating around a novel circular track (the *loop closure* task; Fig. 5). Completion of the first lap produces positive prediction errors between the sharply peaked input from feature inputs learned at the beginning of the trial, and the agent location estimate which is uncertain given the accumulation of PI noise (Fig. 5B).

Decrease in structural error (the difference between the place field separations and their encoded separations on the grid cell sheet) following *online+offline* inference was markedly larger than following *online* learning alone (Fig. 5E). The inferential power of this ‘one-shot’ learning process derives from consideration of the full covariance structure of the feature locations (captured by the CA3 connection weights between place cells), compared to the purely local learning occurring *online*. The system was subsequently able to navigate with dramatically reduced error (Fig. 5Aiv), eliminating prediction errors on subsequent lap completions (Fig. 5B; Supplementary Video 1).

### Coordinated grid-place cell replay as structured information propagation

The *scheduling* of updates in belief propagation is important because messages that do not change the beliefs of neighbours are redundant (Fig. 6A). We scheduled only the place cell whose belief had changed most to broadcast a new message on each cycle (Fig. 6A; see Methods). This *max-update* scheduling was more efficient than simple synchronous schemes, converging with fewer messages (Fig. 6C; see Ref.^49^).

**Figure 6.**
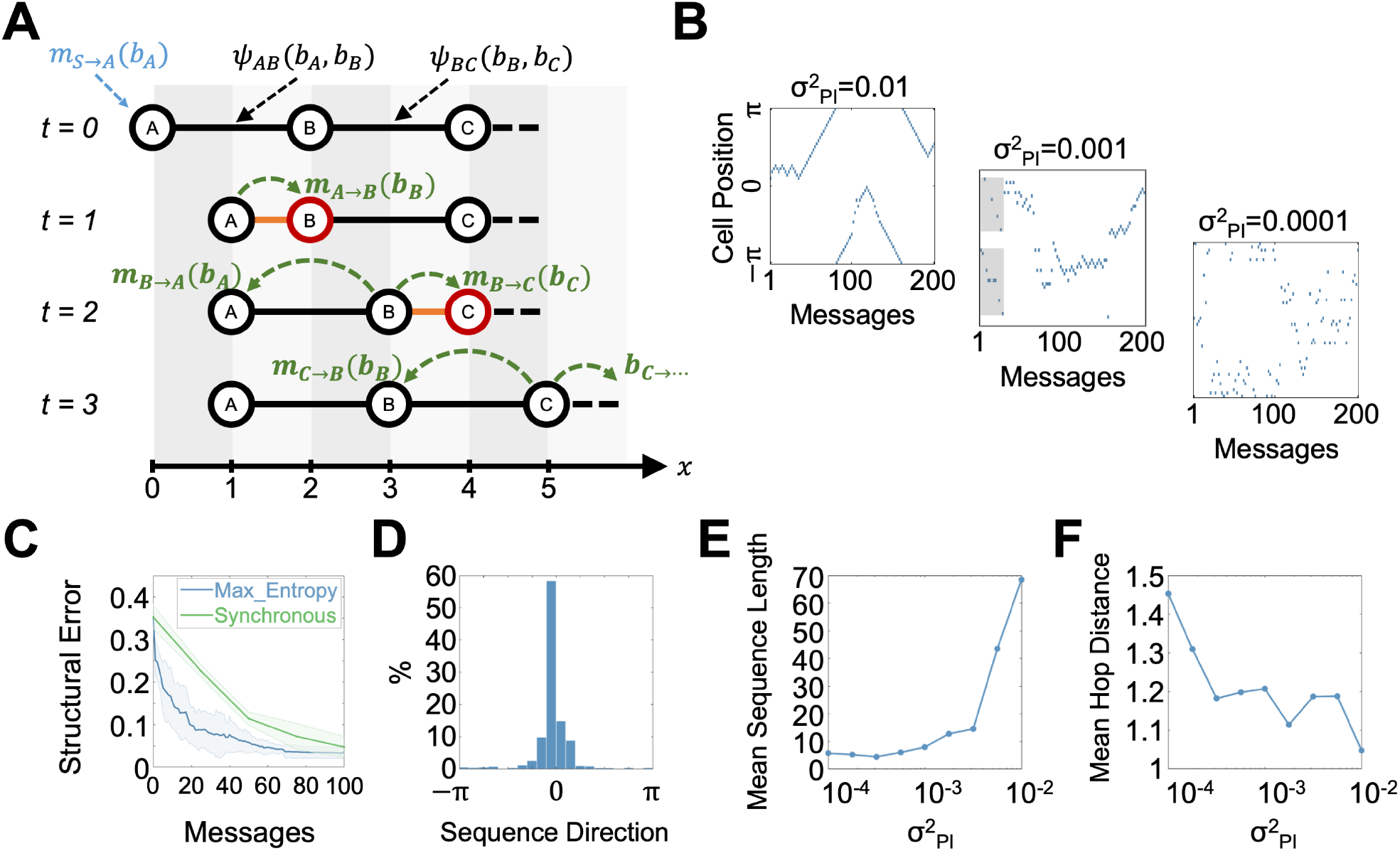
Principled message scheduling during *offline* inference generates sequences of place cell activity. **A)** Place cells (i.e. graph nodes representing conjunctions of environmental features) are connected via their pairwise potentials (*ψ_AB_, ψ_BC_*), which penalize the mismatch between *associative* and *metric* pairwise distances (*δ_ij_*, and *d_ij_*, respectively). (*t* = 0) Environmental sensory input (*m_s→A_*(*b_A_*) causes an update to the belief of place cell A (i.e. updating its synaptic weights to the grid sheet) by making it fire in a new location *b_A_* = 1. (*t* = 1) Place cell A sends a message to B expressing its belief over the location of B, given its own (new) location and the *associative* distance *δ_AB_*, causing B to update its belief. (*t* = 2) Messages from B to A and C only cause C to update its belief, so only C broadcasts at the next time-step. **B)** Examples of PC reactivation sequences in loopclosure task for different values of path integration noise 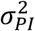 (and therefore pairwise measurement confidence, since 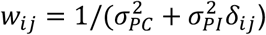; where 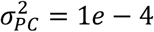). Multiple local sequences can occur in interleaved fashion (Middle, grey shading) and become longer and smoother when pairwise measurements are less confident (Right; also **E, F**). **C)** The ‘MaxEntropy’ schedule (i.e., only the place cell with max entropy change fires in the next-step) converges faster than when all PCs broadcast messages at each time-step. **D)** Forward and reverse sequences occurred approximately equally often.

The sequences of place cells broadcasting messages during *offline* inference in the loop-closure simulation have significant structure (6B). They tend to initially propagate backwards along the track from the animal’s current position, resembling the characteristic reverse hippocampal replay following reward^17^ (Fig. 6B), but also occasionally hop to new locations where remote sequences are initiated^50^ (Fig. 6B,F). These subsequent sequences showed an approximately equal distribution of forward/reverse sweeps (Fig. 6D; see Methods; Supplementary Video 1).

Thus, hippocampal ‘replay’ may reflect correction of local regions of the cognitive graph given new or ‘surprising’ information, as opposed to simple recapitulation of experience^51^. Sequences selectively affect place cells whose beliefs are structurally affected, and terminate when this is no longer the case, ‘hopping’ to remote regions. This leads to smooth sequences in un-converged graphs (novel environments) and more hoppy sequences with experience, where converged regions may be skipped (Fig. 6F). These ‘hops’ marked the separation of ‘replay’ events into distinct sub-sequences (see Methods). Multiple trajectories may also be played out in parallel (e.g. two trajectories alternating under max-scheduling; Fig. 6B, middle, grey shading).

### A neural model of coordinated place cell – grid cell replay

How might belief propagation for *offline* inference be implemented in spikes fired by place and grid cells during replay? We propose a schematic model with a focus on function rather than biological detail (e.g. our ‘place cells’ combine the recurrent connections of CA3 with the connections to mEC of CA1). In the model, minimizing prediction errors between *associative* and *metric* generative models corresponds to synchronizing the propagation of activity through CA3 and mEC, respectively (Fig. 7; see Supplementary Methods; Supplementary Video 2). A ‘message’ is initiated by a place cell spike, which propagates in CA3 via the Hebbian recurrent connections that encode place field separations. In parallel, the same spike initiates activity at the corresponding location on the grid cell module, which then propagates on the grid sheet as a traveling wave, using the same circuitry as path integration in the *online* model and propagating at the same speed as spikes in CA3 (see Methods). Hebbian-like learning strengthens connections from place cells to grid cell which simultaneously receive input in CA3 and EC respectively (Fig. 7A), approximating the algorithmic message-passing implementation (Fig. 7B, C). Firing of the broadcasting place cell is triggered by changes in its synaptic weights to the grid cell population, reflecting correction of the observation model in response to prediction error with the transition model (see Supplementary Methods).

**Figure 7.**
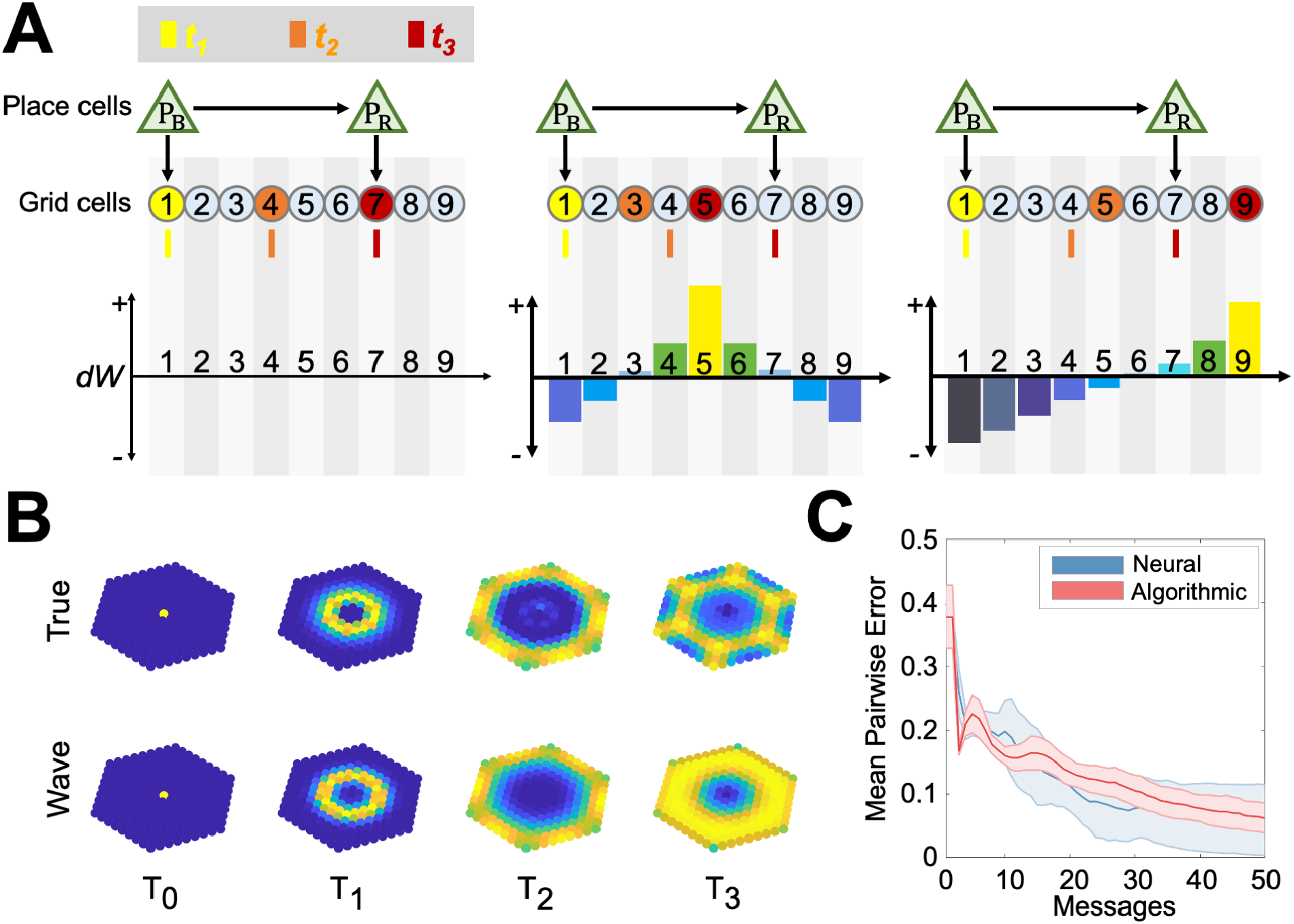
Schematic neural mechanism for probabilistic message passing (see also Supplementary Video 2). **A)** Illustration in 1D. The broadcasting place cell P_B_ sends a spike to receiving place cell P_R_ via recurrent connections in CA3, and also initiates a travelling wave at the corresponding location on the grid cell module via their connections there. (Left) No new learning occurs when the spike and travelling wave arrive at P_R_ and its corresponding grid location (GC7) at the same time, as P_R_-GC_7_ connection will already be strong. (Middle) If the CA3 spike arrives at P_R_ ahead of the travelling wave reaching GC7, the synaptic associations of P_R_ are adjusted towards the currently active GC5 (updating the belief, by increasing P_R_-GC_5_ and decreasing P_R_-GC_7_, see *dW*, below). (Right) Same as middle, except that GC wave reaches GC7 before the CA3 spike reaches P_R_. **B)** Propagating messages as travelling waves in mEC. A neural simulation of travelling waves with a modified Laplacian diffusion kernel (Wave) closely approximates the probabilistic propagation of activity (True), reflecting the accumulation of self-motion noise in the broadening of the wave front. **C)** Comparison of the algorithmic and neural belief propagation performance.

## Discussion

Building on previous work^11,13,15^, we argue that the mEC-HPC system performs spatial inference in two distinct regimes. Given a known ‘cognitive map’ (mapping sensory information to metric space), probabilistic integration allows optimal estimation of current location by *online* combination of uncertainty-weighted self-motion and environmental observations provided by *transition* and *observation* models respectively (Fig. 1). Where these estimates deviate strongly, prediction errors (Fig. 1G) trigger *offline* inference events (Fig. 5B), which propagate local environmental input to remote but structurally associated states, producing coordinated (often sequential) reactivations in place and grid cells (Fig. 6, 7). The effect of *offline* inference is to produce a 2D embedding of the sensory information provided through the place cells, which may facilitate planning or generalization. Although not modelled here, back-projections from grid to place cells, reflecting the metric embedding of their place fields, might therefore also reduce uncertainty in place cells’ firing, producing increased spatial stability in their fields, as observed to occur during sleep^12^.

Partial rescaling of grid patterns^7,21^ and differential shifting of grid and place fields^23^ in response to manipulations of environment sensory input can be understood as joint optimization of *transition* and *observation* models, balancing model priors with new observations. Where prediction errors persist, direction dependent grid pattern shifts may emerge as a result of probabilistic integration of these conflicting cues^7,9^ (whereas boundary-dependent resetting^9^ produces larger shifts than experimentally observed and no rescaling; Fig. 2D, E).

We show that observed grid pattern distortions can be mechanistically linked to inhomogeneity in the sampling or neural representation of the environment^25,26,29^ (Figs. 4A, S3), which might be reflected in behaviour^52^. Thus variation in the confidence, sampling or discriminability of sensory states will produce local changes in grid scale, inducing non-Euclidean structure in the *metric* representation of space (Fig. 1D, 4B). Our model also shows that distortions appear gradually with experience^29^, as the learned mapping from sensory features to metric space (the *observation* model) becomes more confident relative to the estimate of location from path integration (the *transition* model). Given initial learning, *online* localization errors (Fig. S3) should occur immediately following subsequent manipulations to the environmental sensory input, whereas *offline* changes may occur over longer timescales and correlate with replay of the manipulated states (Figs. 4,6; consistent with grid, but not place fields reorganizing significantly during sleep^53^). However, although large prediction errors will cause more easily detectable *offline* inference events, *offline* learning may occur continuously and not necessarily reactivate distinct previously experienced spatial trajectories^51^. We note that strong *associative* connectivity may also contribute to pattern completion, making the place cell representation robust to cue removal^54^.

Theoretical studies have demonstrated how the connectivity of the mEC *metric* space might emerge from a low-dimensional embedding of sensory stimuli^55^, predictive states^35^ or from unsupervised learning during navigational tasks^56^. A crucial difference in our model is that perceptually similar but physically separated compartments will be represented distinctly^57^, reflecting the vectorial translation between them in the transition model (i.e., not simply reflecting the topological state transition structure^35^). Another recent model showed that grid cell like responses can emerge from learning the transition model that best predicts observed sensory stimuli^36^. We instead assume a fixed transition structure but with a variable linear gain, consistent with continuous attractor models^31^ where translation of activity on the grid cell sheet is driven by cells with velocity-dependent firing rates^38,39^. Indeed, a recent study showed that velocity dependence in mEC firing is tied to environmental manipulations^39^.

We propose that *offline* structural inference events correspond to coordinated HPC/mEC replay^16–19,58–60^, which can be viewed as synchronizing predictions from *associative* (CA3) and *metric* (mEC) generative models (Fig. S7). In this way, structural changes to an environment can be propagated to non-local regions of the *metric* embedding, in contrast to models in which these states need to be physically revisited^13^, consistent with the observation that replays do not necessarily repeat experienced trajectories^51^. Prediction errors between the two models may trigger replay events and corresponding sharp-wave ripples^61,62^. To our knowledge, this is the first functional model of coordinated place cell-grid cell replay^18^ (although cf. Ref. ^63^), and provides an alternative to reward-based theories^64,65^ (we note that rewards may themselves represent salient sensory features, independent of their reward value).

Our model makes a number of experimentally testable predictions. Firstly, systematic manipulation of the discriminability of sensory cues distributed within an environment should produce predictable distortions to the grid pattern, observed with increasing experience of an environment. Secondly, replay should be more frequent after structural changes such as shortcuts, blockages or gain manipulations as in the experimental setup of Fig. 5. Thirdly, replay events triggered by specific unexpected sensory observations should become less frequent (Fig. 5B) and smooth (Fig. 6E, F) with continued experience, if the observations remain stable. Fourthly, multiple local replay events may occur in inter-leaved fashion (Fig. 6C, *middle*, grey shading). Fifthly, we predict the existence of travelling waves in grid cells (as a function of their spatial phase; see also^66,67^, Fig. 7). Lastly, initial messages propagating from the animal’s current location may not cause subsequent messages in remote regions of the graph which are already sufficiently converged (messages will not cause changes in the beliefs of their recipients), although activity in mEC will continue to propagate. Thus grid cell replay could thus be detectable in the absence of simultaneous place cell replay^19^ (but place cell replay requires the grid cell *transition* model and so depends on mEC^68^).

Our proposed structure learning framework can account for diverse phenomena observed in the HPC-mEC system, and makes several novel, experimentally testable predictions.

## Methods

### *Online* recursive Bayesian estimation

The transition matrix *T* defines the probability of transitioning from agent location ***x***’ to location ***x***, and is a function of the perceived current velocity ***û*** and *transition* model gain ***α*** = [*α_x_*, 0; 0, *α_y_*]. Since our *metric* space is periodic, *T* accounts for cyclic transitions ***c**_mn_*, with Gaussian noise proportional to the perceived velocity ***Ϻ~u*** + *N*(0, diag(***u***)**Σ**_PI_):

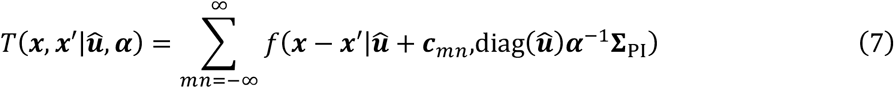

where *f*(***x***|***μ*, ∑**) is a multivariate Gaussian PDF, ***c**_mn_* = 2***α***^1^(*mv*_1_ + *nv*_2_) and 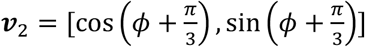 define the unit vectors of a hexagonal lattice^69^ with grid pattern orientation *ϕ* and diag(·) produces a diagonal matrix from a vector input. Since most of the mass is associated with shorter transitions, in practice we approximate the full distribution with a finite number of periodic summations (i.e. ignore the tails; 5 cycles in our simulations; Fig. S1B).

The *observation* model defines the likelihood of the current environmental sensory inputs (i.e. the population vector of place cell firing ***P***, where 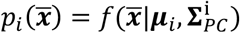 is the firing rate distribution of place cell *i* over physical space 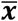) given the predicted *metric* location ***x***, via a thresholded weighted sum: *H*(***P|x***) = [∑_*i*_ *B_i_*(*x*)*P_i_*]^+^. Here, *B_i_*(*x*) is the location distribution of landmark *i* in *metric* space, which would be encoded biophysically in the learned [*N_P_* × *N_G_*] matrix ***B*** of synaptic weights from place to grid cells (i.e. the *g^th^* row and *i^th^* column of *B* is the distribution ***B**_i_*(·) evaluated at the location of the *g^th^* grid cell). The normalization constant *K* = ∫ *G*(***x***)*d**x*** in Eq. 2 simply sums over the current grid cell activity and might biophysically be implemented by inhibitory interneurons.

### *Online* learning of structural priors

In the *online* model, the place-grid cell weight matrix ***B*** is learned using the BCM rule:

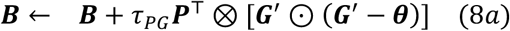

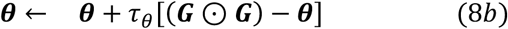

where *τ_PG_* = 1*e* – 4 is the learning rate and ***G***’ and ***H*** are column vectors whose elements are the *transition* and *observation* models estimated at the locations of a finite set of grid cell locations. ⊙ is the element-wise (Hadamard) product between two vectors and ⊗ is their outer product. The sliding threshold *θ* ∈ *R*^1×*N_G_*^ provides adaptive synaptic normalization, where *τ_θ_* ≈ 10*τ_PG_*. Learning takes place between the *apriori* distribution ***G***’ and the current sensory observation ***H*** (i.e. before the observation correction to ***G***’).

### The *offline* probabilistic graphical model

The pairwise potentials 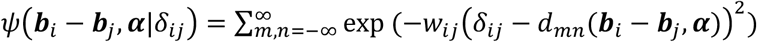 penalize differences in the pairwise distances encoded by *association δ_ij_*, and those that would be computed by comparing their absolute encodings in *metric* space *d_mn_*(·); i.e. they encourage a metric embedding that reflects the associative distance. The *metric* distance function 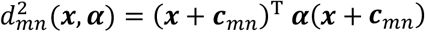 defines the pairwise distance between the encoding of locations *i*. and *j* in mEC in *metric* space, and is dependent on the gain factor ***α***. of the *transition model.* Pairwise measurements are assumed to have confidence *w_ij_*, (inverse variance) that increases with decreasing inferred distance (i.e. *w_ij_*, = 1/(*σ_PC_* + *σ_PI_δ_ij_*)). The *transition model* gain is assumed to have a Gaussian prior *p*(*α*) = exp (–*w_**α**_* (***α*** — ***α***_0_)^T^(***α*** – ***α***_0_)), the *w_**α**_* term representing the confidence in the prior gain value ***α***_0_. The periodic offset term *c_mn_* is the same as defined for the transition model.

### Associative encoding in the hippocampus

The associative distances are recovered from the [*N_P_* × *N_p_*] synaptic weights in CA3 ***A***. Under a random-walk behavioural trajectory, the simple modified Hebbian learning rule:

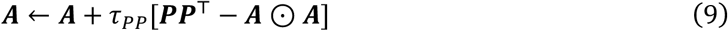

where ⊙ is the element-wise (Hadamard) product and *τ_PP_* the learning rate. The synaptic weights can be shown to converge to 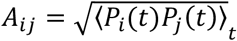, the square-root of the correlation between the firing rates of two PCs^70^. Where place fields have uniform receptive field widths (*σ_PC_*) and peak firing rates, the Euclidean distance between place fields *i* and *j* can be inferred via the simple transformation^46^:

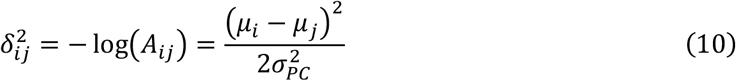

The recovered distance is therefore scaled by the receptive fields’ variance (the Bhattacharyya distance^46^), and so relates to ‘discriminability’ (Fig. 1D). CA3 synapses effectively average over multiple pairwise measurements. By assuming that noise in the pairwise distance measurements scale linearly with distance, both the mean and variance of the Gaussian describing this distribution is efficiently encoded in a single PC-PC synapse.

### Simplified analysis of the probabilistic graphical model likelihood

To characterize model predictions in the environmental rescaling^71^ and gain change^7^ experiments, we studied a reduced version of the full graphical model (Eq. 3; see Supplementary Methods for full derivation). In 1D, given a linear *observation model x* = *H*(*x*’) = *K*_1_ + *Kx*’ and a large number of evenly spaced place fields, Eq. 3 simplifies to (see Supplementary Methods):

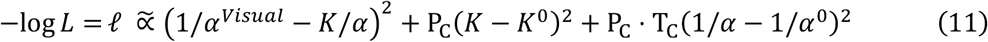

which can be solved analytically. A similar reduction was applied to the 2D case when considering differential shifts in grid and place fields^23^.

### Belief propagation for *offline* inference

Belief propagation^48^ is an iterative, two-stage local message-passing scheme in which, at each iteration *n*, a feature node (i.e. a place cell) first updates its own belief (connections to the grid cells) 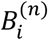 by integrating messages from connected nodes *j* ∈ Γ***i*** with its own prior belief 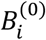 (Fig. 5C):

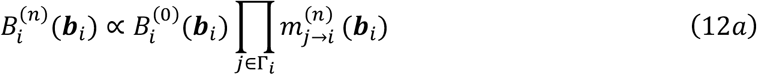

The message from node *j* to node *i* (*m_j→i_*) communicates its belief over the distribution of the locations of place cell *i* in grid cell space, conditioned on its own location distribution:

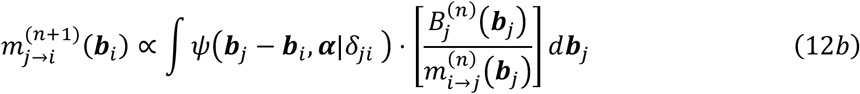

where the pairwise potentials *ψ*(·) are the same as those described in the full likelihood function (Eq. 3). The graph converges when new messages cease to change the beliefs of their recipient nodes.

### Scheduled message passing on the place cell graph

‘Synchronous’ belief propagation computes belief updates for each step before broadcasting all new messages in the next step. In simulations, we demonstrated that scheduling message broadcasts based on internal ‘message tension’ (divergence between previous and updated belief given new messages) produced faster and more accurate convergence (Fig. 6B; see also Elidan et al.^49^). Message tension between the node’s previous 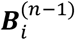 and updated 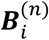 beliefs is defined as:

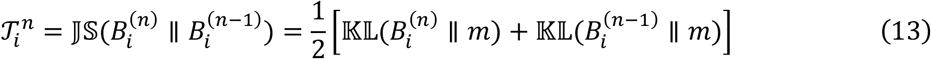

where 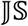 is the Jensen-Shannon (symmetric K-L) divergence, where:

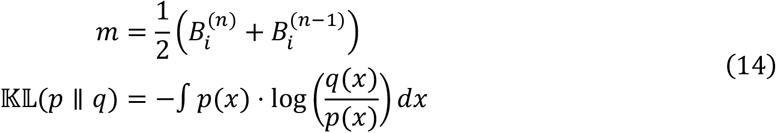

When the message tension is below a pre-defined threshold 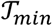, a node has converged and ceases to broadcast new messages. This mechanism is similar to the prediction error between *transition* and *observation* models used to trigger *offline* inference, with the exception that it uses the symmetric divergence measure 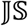 rather than 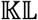.

### Traveling waves in neural media

In simulations, the traveling waves in mEC are simulated explicitly by calculating the true messages conditioned on the sending nodes’ current beliefs at each time-step (Eq. 5B). In the ‘neural model’ (Fig. 7), messages were approximated as waves propagating radially from an initial stimulation on the mEC sheet using a modified mechanical wave model, as used to describe oscillations in water:

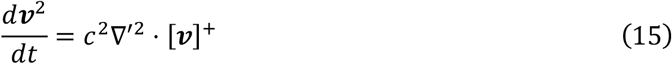

where *c* is the speed of wave propagation, [·]^+^ is a threshold linear activation function, and the modified spatial Laplacian operator ∇’ is a symmetric 2D Gaussian filter with variance equal to the PI noise (see Supplementary Methods for extended discussion).

## Acknowledgements

We thank the groups of Caswell Barry and Francesca Cacucci for use of their data and Martin Pearson (UWE and Bristol Robotics Lab), Dan Bush, Andrej Bicanski and Tim Behrens for valuable theoretical discussions.

We acknowledge funding by the ERC Advanced grant NEUROMEM, the Wellcome Trust and the European Union’s Horizon 2020 research and innovation programme under grant agreement No. 785907 Human Brain Project SGA2. The authors declare no competing financial interests.

---

**Figure S1.**
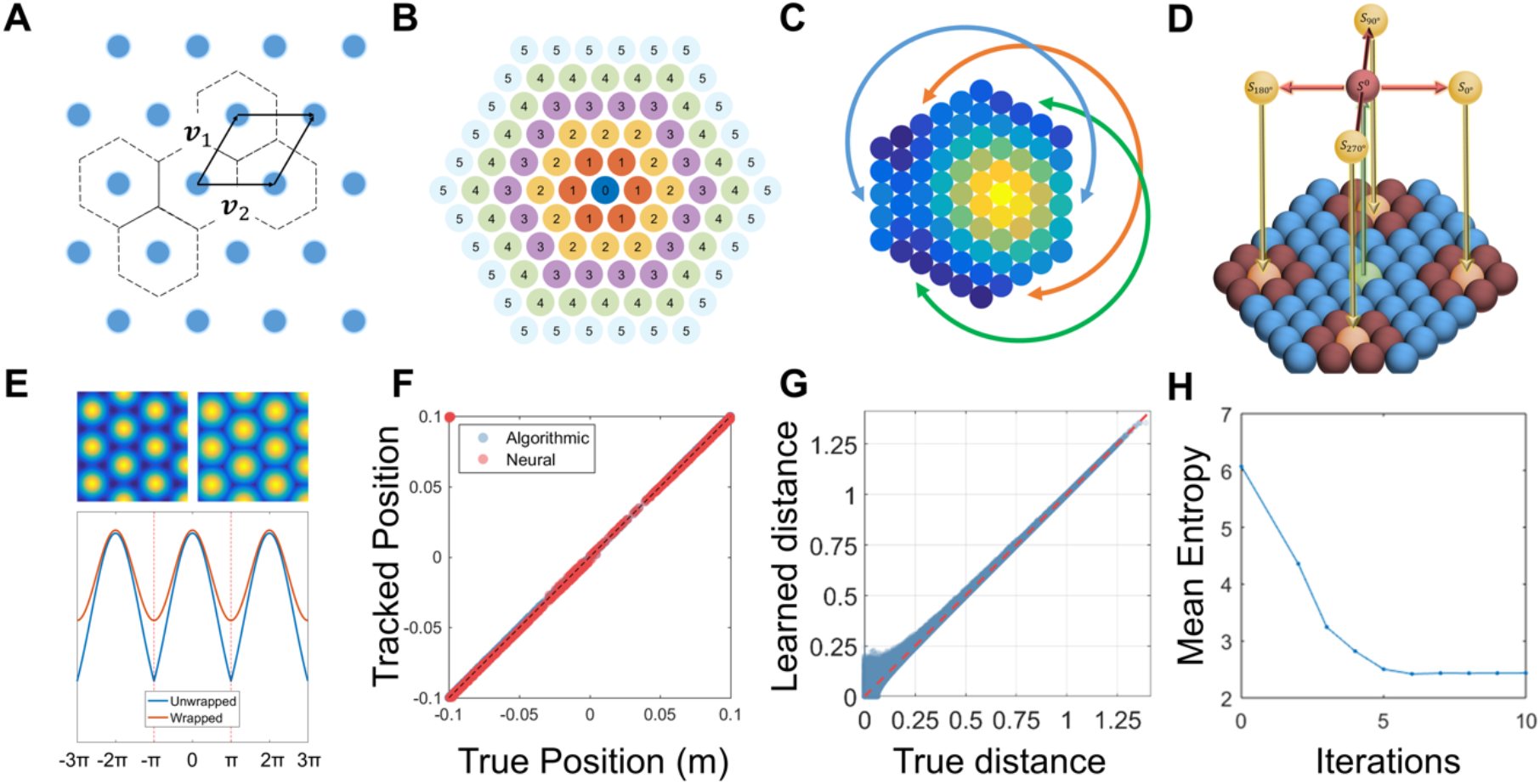
Implementational details of the *online* system. **A** The mapping from real to grid cell space can be considered as the subtraction of a mapping vector [*i·V*_1_ +*j*· *v*_2_], where each grid field can be described as a point on a 2D lattice with basis vectors *v*_1_ and *v*_2_. Each grid field has associated with it a Voronoi domain, defined as the region within which any point is closest to the corresponding grid field. When mapping from grid-to-real space, the vector of the closest grid field is subtracted. **B** The wrapped Normal distribution is a summation of the likelihoods of the current position estimate being at any one of an infinite number of periodic tilings (here, five wrappings are shown). **C** The grid cells are connected periodically to produce a ‘twisted-torus’ topology. **D** Illustration of the shifter cell mechanism. Each ‘readout’ grid cell is connected to four ‘shifter’ grid cells and a single self-connecting cell. **E** Illustration of the difference between wrapped and non-wrapped Gaussian distributions. **F** Correspondence between the neural shifter cell mechanism and the algorithmic transition function. **G** Pairwise distances between place fields can be inferred from the strengths of Hebbian connections. **H** Mean entropy in the beliefs of each place cell over their encoded location in grid space (encoded in place-grid cell connection weight distributions decreases with iterations during *offline* inference.

**Figure S2.**
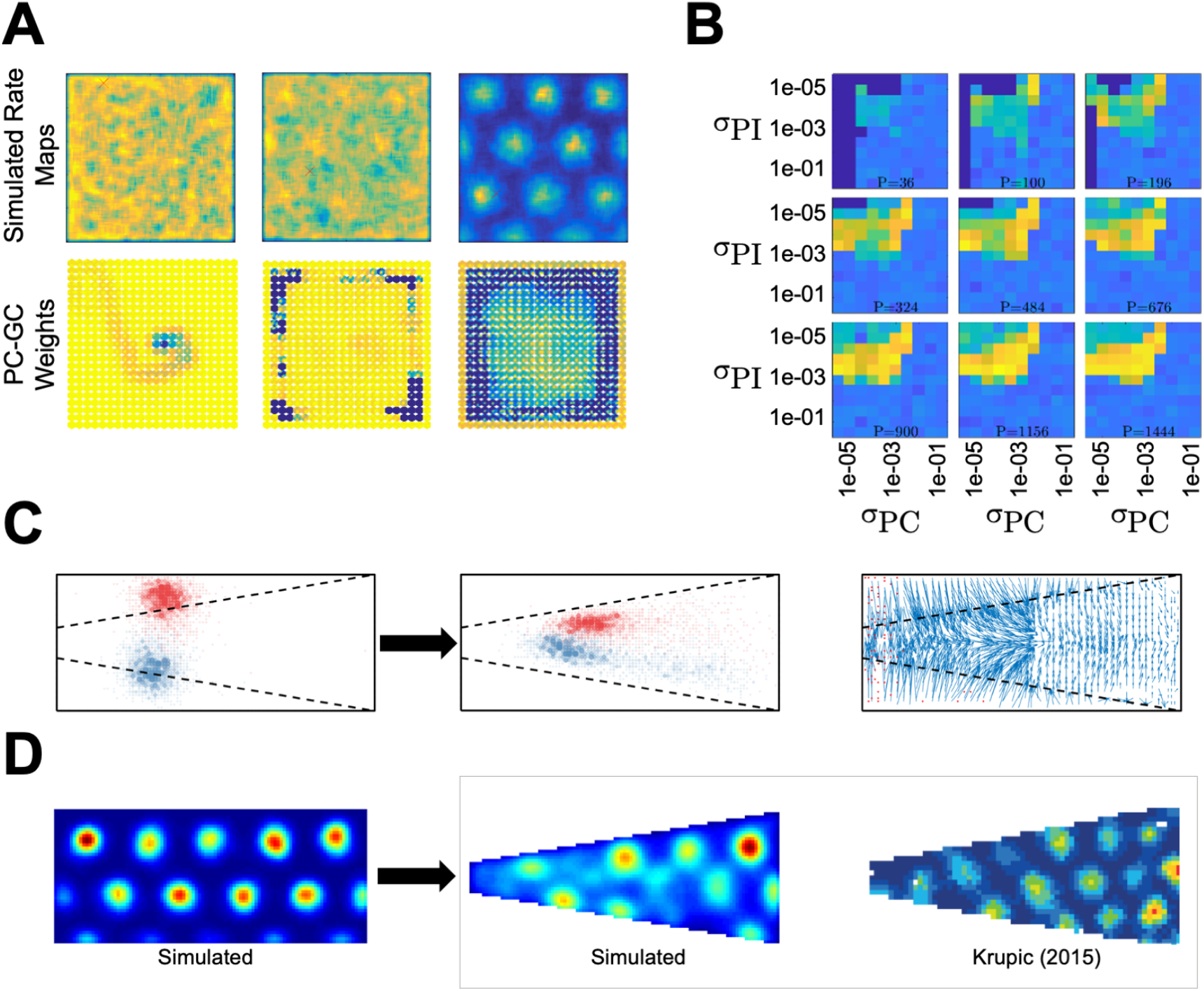
**A** Convergence of grid patterns and place-grid cell weights over three periods (columns). The place-grid cell weights *(bottom row,* colour denotes normalized connection strength) converge fastest near to the boundaries (as seen in development; Muessig et al., 2015) and corners of an environment, since trajectories through are more directionally constrained. Top row shows corresponding grid patterns (colour denotes normalized firing rate). **B** Convergence of stable grid patterns over sensory noise (*σ_PC_*) and path integration noise (*σ_PI_*). Colour denotes grid score (Sargolini et al., 2006). *P* indicates number of place cells used in simulations. **C-D** Environmental deformations caused by prior structural beliefs. **C** Place fields generated by the boundary vector cell model (Hartley et al., 2000). An animal’s perception of a trapezoidal environment *(right)* may be influenced by place-BVC associations learned in a previous rectangular environment *(left).* In the trapezoid, simulated place fields shift with the wall (**D**), causing similar distortion of the grid pattern.

**Figure S3.**
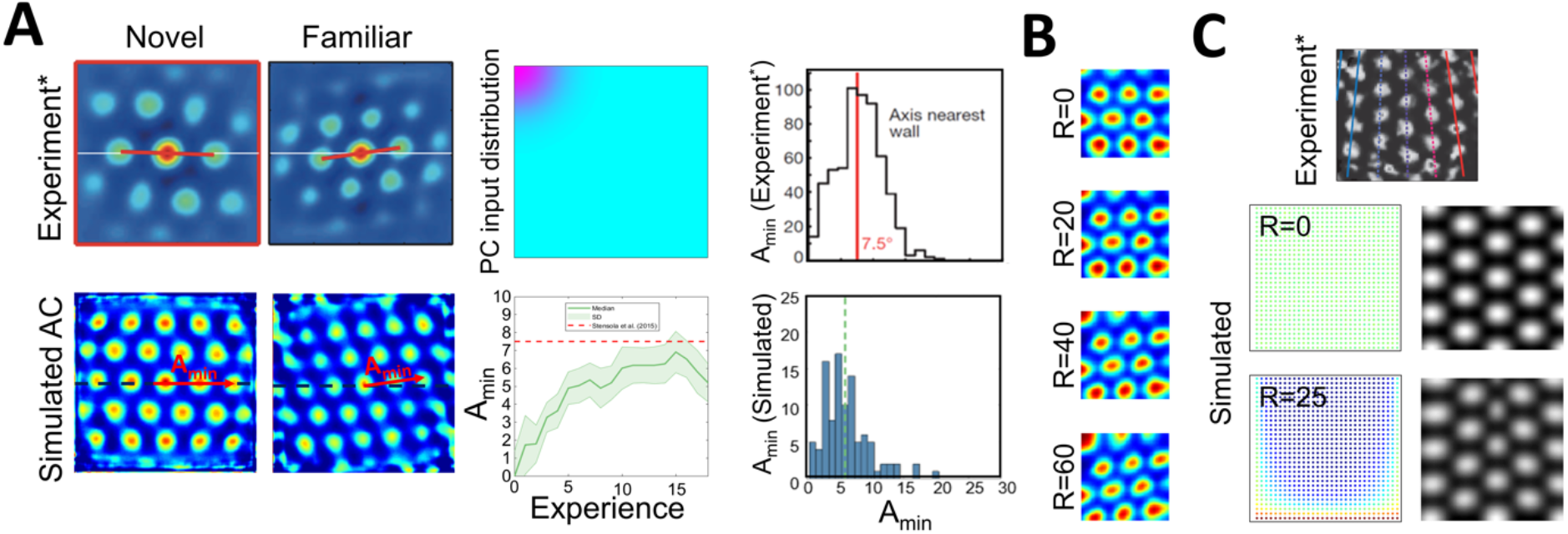
Grid distortions due to inhomogeneous environmental inputs in the *online* model. **A)** When environmental inputs are concentrated in one corner of the environment *(top-middle panel),* the resultant grid cell firing rate maps undergo a shearing distortion which produces an orientation offset (A_min_; *bottom-left panels).* This offset increases with experience, as grid cell firing becomes increasingly dependent on the maturing sensory inputs *(bottom-middle panel),* matching experiments in which rats always entered the box at the same corner *(top-left panels;* Stensola et al., 2015) The size of the experimental and simulated offsets are similar *(right panels).* **B)** Simulated distortions based on an exponential decay in place cell input from one corner as a function of the decay parameter (R). **C)** Concentrated place field input along one wall and both corners *(bottom left panel)* causes another distortion pattern *(bottom right panel)* also observed experimentally (*top* adapted from Stensola et al., 2015).

**Figure S4.**
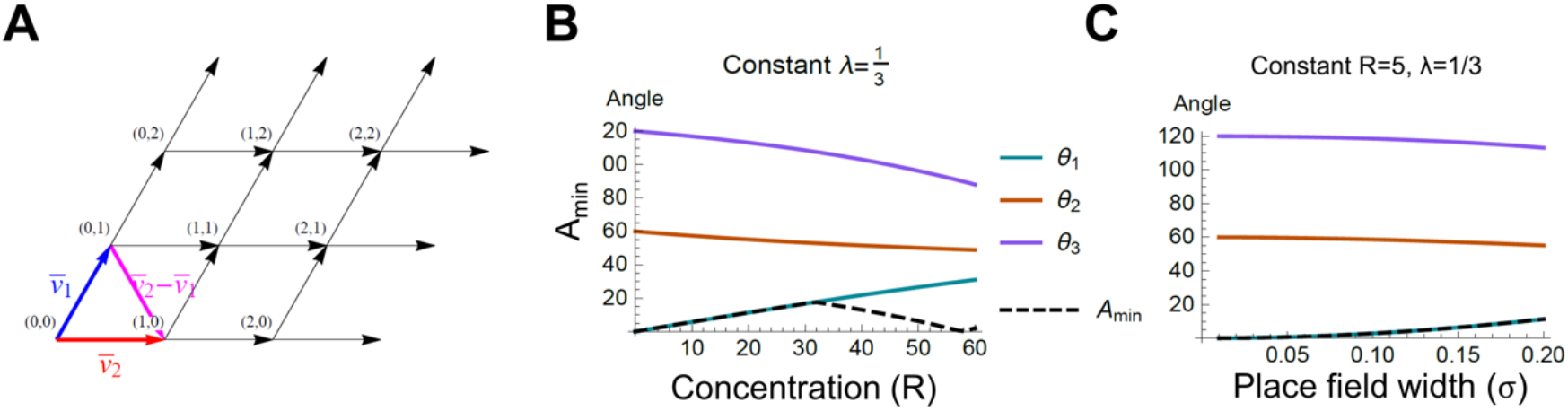
Mathematical analysis of shearing of the grid pattern due to inhomogeneous environmental inputs. **A** The orientation resulting from the shearing operation can be calculated by analysing the angles of the sheared hexagonal lattice describing the centroids of the grid fields. **B, C** The analysis predicts that the orientation offset should be dependent on both the strength of the place fields’ density / firing rate imbalance *R* (see Supplementary Methods 1.7 and Fig. 2) (**B**) and their tuning widths (**C**).

**Figure S5.**
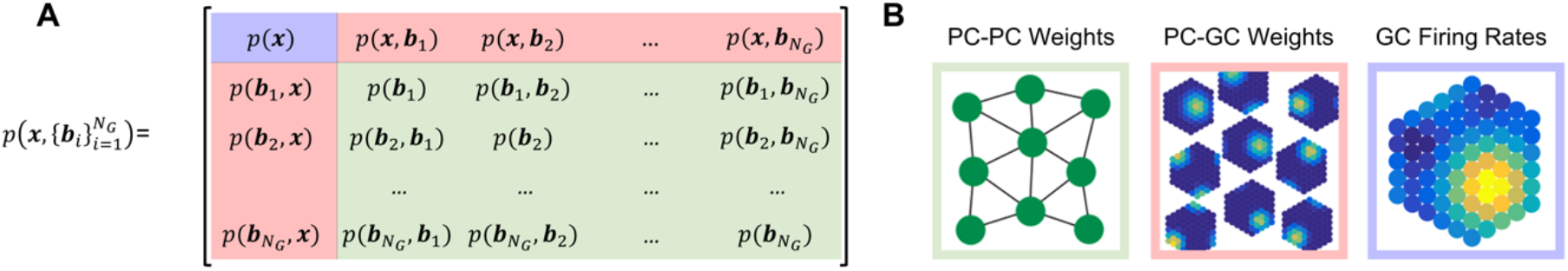
Anatomy of a SLAM system. The joint location-map probability distribution **(A)** is represented in the firing rates and synaptic weights within the HPB-mEC system **(B).**

**Figure S6.**
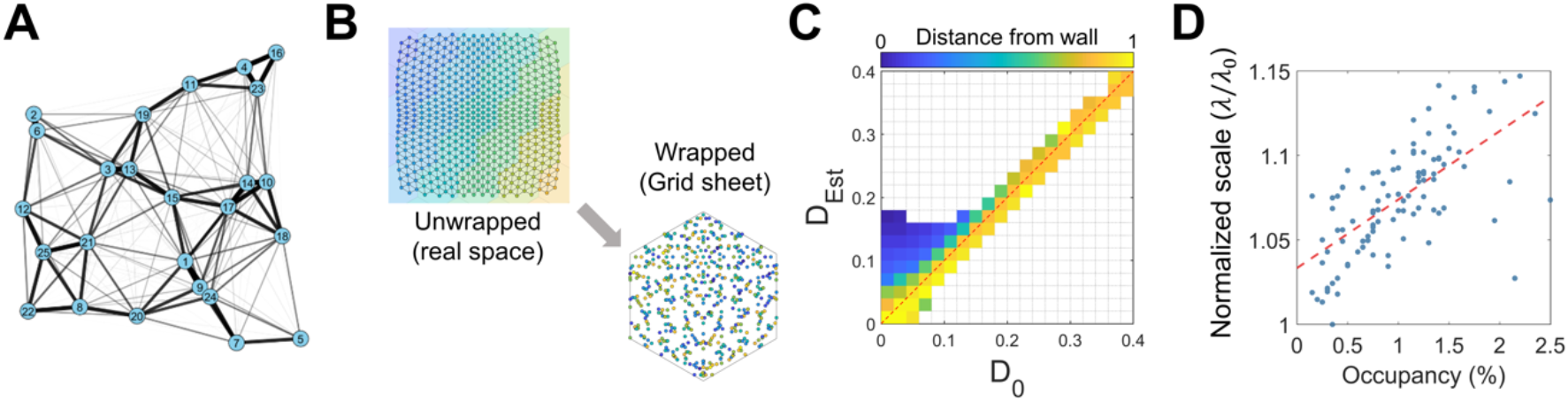
**A** Spring network analogy of the associative structure of an environment. Edge ‘stiffness’ is inversely proportional to the variance in the Gaussian observation. **B** ‘Wrapping’ physical space to encoded location on the grid sheet. Each colour indicates the tiling of the based grid sheet’s domain in real space. **C** Pairwise distances near the edges of the environment are overestimated due to under-sampling when the agent preferentially explores the middle of an environment. Colours denote the distance of the pair of PCs *i* and *j* from the walls of the 1×1 m^2^ environment 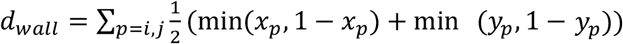. **D** Resulting local scale is proportional to the occupancy. C When the grid scale is smaller than the size of structure being encoded, we can think of ‘wrapping’ the structure onto the grid sheet. Here, colours denoted different tilings of the base metric tile (the Voronoi region of a given grid cell).

**Figure S7.**
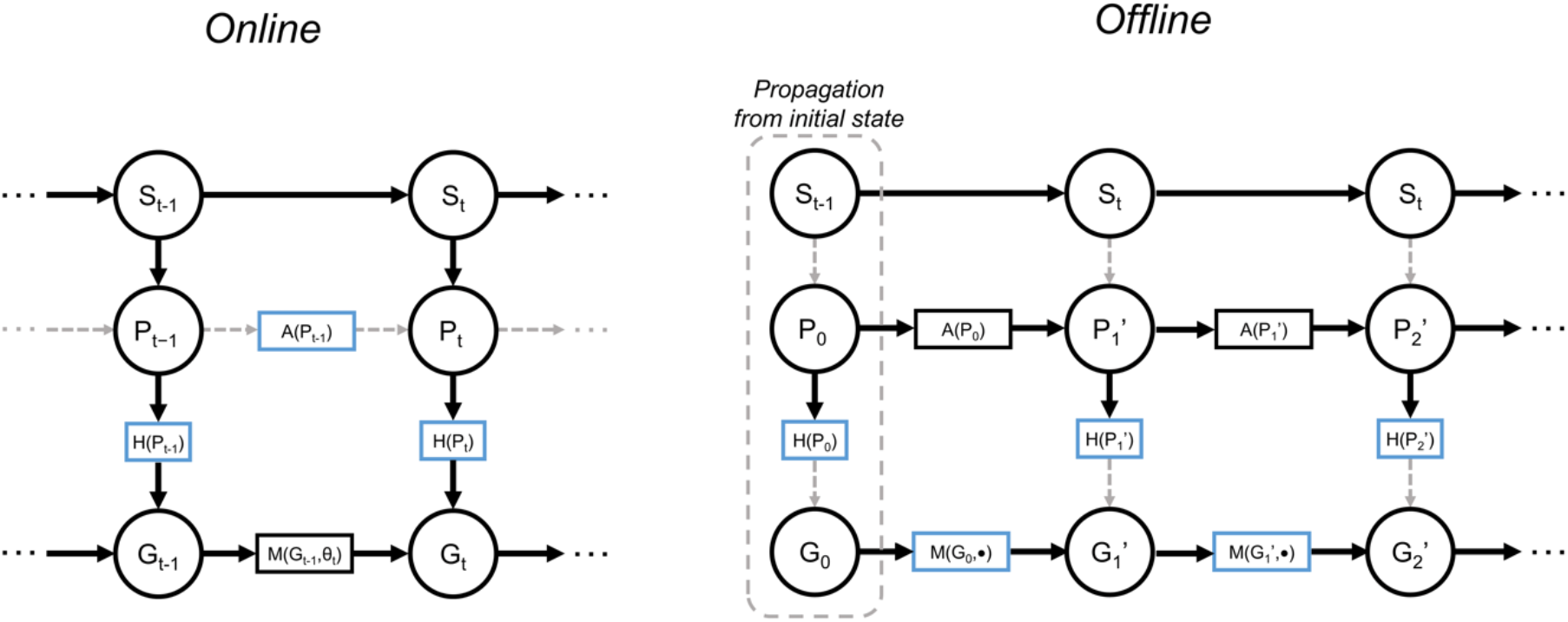
An alternative view of the *online* and *offline* models. Place cells *P* are driven by real-world stimulus S. During *online* exploration, the *associative* generative model is learned, but does not generate predictions. During *offline* inference, the *metric* generative model is corrected towards the predictions being generated by the *online* model, which becomes a surrogate for sensory stimuli as would be generated by the real world model.

**Table S1.** Algorithm detailing the overall dual-systems hypothesis of *online* and *offline* localization and learning in the HPC/mEC.

| **Algorithm:** *Online* localization and mapping with prediction error initiated *offline* inference |  |
| --- | --- |
| 1: initialize: $\mathbf{A} \sim \mathcal{U}(0, \Delta)$, $\mathbf{B} \sim \mathcal{U}(0, \Delta)$ | % Initialize weights to small values $\sim \Delta$ |
| 2: while $k < K$ do: | % For the duration of the simulation |
| 3: do_movement_update(); | % Update state via path integration |
| 4: make_observation(); | % Compute PC firing |
| 5: if $\epsilon_k > \epsilon_0$ do: | % If prediction error, do *offline* inference |
| 6: while any $\mathcal{T}_i < \mathcal{T}_{min}$ do: | % Loop until cells below tension threshold |
| 7: $u = \text{argmin}(\mathcal{T})$; | % Find node with max. message tension |
| 8: compute_and_broadcast_message(u); | % Broadcast message to neighbours of $u$ |
| 9: for $t \in \text{Neighbours}(u)$ do: | % Loop over neighbours of $u$ |
| 10: update_belief($t$); | % Update belief of node $t$ |
| 11: update_message_tensions($t$); | % Compute change in belief of node $t$ |
| 12: update_PC_GC_weights(); | % Do *associative* to *metric* map learning |
| 13: update_PC_PC_weights(); | % Update *associative* weights |
| 14: do_measurement_update(); | % Correct state estimate |

