## Supplementary Materials for "Replay as structural inference in the hippocampal-entorhinal system"

### Supplementary Information

#### 1 Agent simulations

The *online* model simulated an agent navigating either randomly in an open field, or clockwise around a circular track. Inferred *offline* structure was computed numerically/analytically from the probabilistic graphical model (Main Text Eq. 3) or by taking the maximum of each landmark's encoded distribution following message passing.

##### 1.1 Online model

The firing pattern of a given GC (Fig. 1; Supp. Fig. 3; Main Text Fig. 1A) is generated by plotting the activity of the GC against the location of the simulated agent in physical space. If there is a preserved 'bump' of activity on the GC sheet (in metric space), as this bump moves according to path integration a given GC will periodically become active and inactive. Thus, a larger grid scale corresponds to a smaller velocity on the GC sheet; conversely, a larger velocity will cause the activity bump to complete more 'laps' of the sheet for a given movement in physical space (more vertices in the readout grid pattern).

##### 1.2 Offline model

Grid cell firing patterns in Figures 2, 3 and 4 were generated by directly projecting weighted place cell activity across space (i.e. the solution corresponds to the structure embedded via the *observation* model, and is not affected by *online* path integration), where the locations of their respective place fields are obtained from the maximum-likelihood solution to the simplified objective function detailed in Section 4.

Note that pairwise associative distance is inversely related to the scale of the grid pattern readout, since travelling between two place fields with a small associative distance will mean travelling less far along the mEC metric space; thus, there will be fewer 'revolutions' of this periodic space for a given unit of movement in physical space, producing a large grid scale (Fig. S6B).

#### 2 Localization-induced distortions during *online* exploration

Grid firing patterns exhibit global shear-like distortions[28] and 7.5-8° offsets of one of the grid axes[28,42] to the walls of square environments. In Stensola et al.[28], rats were introduced into the same corner of the box at the start of each trial. In Butler et al.[24], shearing developed following the introduction of reward[24]. In both experiments, shearing developed with increasing experience[24,28]. We hypothesized that these distortions reflect non-uniform environmental inputs to the grid module, either reflecting their natural distribution[24,28] or inhomogeneous sampling of environmental locations[25].

Given a correctly learned map, during *online* localization, increasing the firing rates or connection strengths of place cells with fields in one corner of the box biases the *observation* model estimate, producing a shearing in grid patterns that are initially symmetrical and aligned to a box wall, and an alignment offset relative to the wall matching experimentally observed values (Fig. S3A, right). Other spatially biased distributions of place cell inputs produce other observed grid distortions (Fig. S3C). These distortions arise as the combined estimate of location becomes biased towards over-represented sensory inputs, and develops with experience as the place-grid cell connections strengthen over time, increasing reliance on *observation* over *transition* model estimates (Fig. S3A, Supplementary Methods). This would also cause grid distortions following the introduction of a rewarded location[24,43], assuming that this causes inhomogeneous place cell inputs[26,44].

#### 3 Generalized shifter cell mechanism

To implement the movement update in a neurally plausible way, we adapted the 'shifter cell' model[3]. We assume two populations of grid cells, the first being the 'readout' neurons to which we have referred so far and produce the simulated grid patterns, and a second 'shifter cell' layer which are driven by and back-project to the readout cells.

In the original model, the shifter cells back-project to spatially adjacent readout cells to those from which they are driven. In our model, these back-projections (Fig. S1D) have Gaussian tuning with a variance $\boldsymbol{\Sigma}_{PI}$, such that given physical movement in the absence of corrective sensory input, the movement distribution would broaden over time (Fig. 1A). The back-projections are tuned to a fixed offset distance proportional to $V_{max}$, which is the maximum velocity at which the animal can travel.

In our simulations, we assume that each readout GC has four corresponding shifter cells $S_\theta^i$, each tuned to one of four principal directions $\theta$ separated by 90°. Each shifter cell fires in proportion to the projection of the current heading velocity vector $\hat{\boldsymbol{u}}_t$ on to it preferred firing direction, multiplied by the drive from its corresponding readout cell $i$.

The current architecture would only allow the current pattern of activity to be translated by a fixed amount in an arbitrary direction but not remain stationary, since the offset connectivity has a fixed offset on the grid sheet. Thus, a fifth corresponding shifter cell type is assumed, $S_0^i$, which back-projects only to the corresponding readout GC. These cells are effectively self-connections, however their firing is independent of heading direction and negatively proportional to the current velocity magnitude. The shifter cell firing is then defined as:

$$S_\theta^i(\boldsymbol{v}) = \& \left[ \left( \frac{\boldsymbol{v} \cdot \hat{\boldsymbol{e}}_\theta}{V_{max}} \right) \cdot G_i \right]^+ , \quad \text{for } \theta = [0,90,180,270] \tag{1a}$$

$$S_0^i(\boldsymbol{v}) = \& \left[ \left( \frac{V_{max} - |\boldsymbol{v}|}{V_{max}} \right) \cdot G_i \right]^+ \quad \text{when stationary} \tag{1b}$$

Where $[\cdot]^+$ is a threshold linear activation, such that two shifter cells with preferred directions 180° apart never fire together. $\hat{\boldsymbol{e}}_{\theta_j}$ is a normalized unit vector denoting the preferred firing direction of cell $j$. Note that this notation accounts for both direction and magnitude; the magnitude is scaled as a proportion of the maximum velocity $V_{max}$, which is assumed to be larger than the maximum velocity of the animal as measured per timestep $\Delta t = 1e - 4$. The back-projecting weight matrices are defined as:

$$W_\theta(\boldsymbol{x}_i, \boldsymbol{x}_j) = N_W(\boldsymbol{x}_i - \boldsymbol{x}_j, 0, V_{max}\boldsymbol{\Sigma}_{\text{PI}}) \tag{2a}$$

$$W_0(\boldsymbol{x}_i, \boldsymbol{x}_j) = \delta(\boldsymbol{x}_i, \boldsymbol{x}_j) \tag{2b}$$

Where $\boldsymbol{x}_i$ and $\boldsymbol{x}_j$ are the positions of the readout cells in grid space. Thus, as velocity increases the firing rates of the symmetric (static) and asymmetric shifter cells decrease and increase, respectively. Under this scheme, the effective back-projection from the shifter cells to the readout population is a linear interpolation of the current activity (provided by the static self-connecting cells $S_0^i$) and an asymmetric drive (provided by the cells $S_\theta^i$). Thus, despite the shifter cells having a fixed offset distance, the weighting between the symmetric and asymmetric shifter cells allows an effective back-projection to an arbitrary distance when the velocity is in the range $|\boldsymbol{v}| \in [0, V_{max}]$. The system can then be modelled as a first order ordinary differential equation:

$$\tau \frac{dG_i}{dt} + G_i = S_0^i(\boldsymbol{v}) + \sum_j \sum_{\theta=0:90:270} S_\theta^j(\boldsymbol{v}) \cdot W_\theta(\boldsymbol{x}_i, \boldsymbol{x}_j) \tag{3}$$

Where $G_i$ is the firing rate of GC $i$. Since both the outgoing and recurrent weight matrices to the static shifter cells are diagonal matrices with unity weights, the explicit dependence on the weight matrix is dropped to simplify the notation. Although we do not simulate it explicitly, this mechanism could be readily extended to the case of variable path integration noise by instead assuming a population of speed dependent cells, each with a preferred tuning at some particular velocity, then adding Gaussian noise into the upstream velocity estimator. Thus, for zero noise in the velocity estimate, the activity bump would be translated only by the activity of one sub-population of speed and angle dependent shifter cells.

#### 4 Joint offline optimization of the transition and observation models

We begin with the assumption of a 1D track of length $L$, with $N$ feature locations distributed uniformly along its length. Beginning with the log-likelihood function:

$$l = \sum_{i,j} w_{ij} \left(\delta_{ij} - d_{ij}(\boldsymbol{X})\right)^2 + \sum_i w_i^{Obs} \left(x_i - x_i^0\right)^2 + w^\alpha \left(\frac{1}{\alpha} - \frac{1}{\alpha^0}\right)^2 \tag{4}$$

where $w^{\alpha}$ is the confidence of the *transition* gain prior $\alpha^0$, $w_i^{Obs}$ is the confidence associated with the prior belief of landmark $i$ and $w_{ij}$ the confidence associated with pairwise observation $(i,j)$. We can then substitute in expressions for the *associative* and *metric* pairwise distances:

$$l = \sum_{i,j} w_{ij} \left( \frac{1}{\alpha^{Visual}} \|\bar{x}_i - \bar{x}_j\| - \frac{1}{\alpha} \|x_i - x_j\| \right)^2 + \sum_i w_i^{Obs} \left( x_i - x_i^0 \right)^2 + w^{\alpha} \left( \frac{1}{\alpha} - \frac{1}{\alpha^0} \right)^2 \quad (5)$$

Next, we assume a linear observation model $x = H(\bar{x}) = K_1 + K\bar{x}$, which defines the mapping between physical location on the track $\bar{x}$ and location on the grid sheet $x$, where the prior observation model $H^0 = H(\bar{x}_i^0)$. We assume also that we do not care about optimizing the offset, such that $K_1^0 = K_1$; this is equivalent to assuming that the weights are fixed at the start of the track, as they might be if anchored by strong stimuli. Cancelling terms then gives:

$$l = \sum_{i,j} w_{ij} \left( \frac{1}{\alpha^{Visual}} - \frac{K}{\alpha} \right)^2 \left( \bar{x}_i - \bar{x}_j \right)^2 + \sum_i w_i^{Obs} \bar{x}_i^2 (K - K^0)^2 + w^{\alpha} \left( \frac{1}{\alpha} - \frac{1}{\alpha^0} \right)^2 \quad (6)$$

Assuming that connections only exist between place cells with adjacent feature locations, each pair will then be separated by a fixed distance $\Delta\bar{x}$ and that all priors have equal confidence $w_i = w$:

$$l = 2 \sum_i w \left( \frac{1}{\alpha^{Visual}} - \frac{K}{\alpha} \right)^2 (\Delta\bar{x})^2 + \sum_i w_i^{Obs} \bar{x}_i^2 (K - K^0)^2 + w^{\alpha} \left( \frac{1}{\alpha} - \frac{1}{\alpha^0} \right)^2 \quad (7)$$

Which when N is large is approximately:

$$l \approx 2w \left( \frac{1}{\alpha^{Visual}} - \frac{K}{\alpha} \right)^2 \frac{L^2}{N} + w^{Obs} (K - K^0)^2 \frac{NL^2}{3} + w^{\alpha} \left( \frac{1}{\alpha} - \frac{1}{\alpha^0} \right)^2 \quad (8)$$

Which, diving through by a constant, aggregating terms and defining the prior confidence and transition confidence scores $\mathrm{P_C}$ and $\mathrm{T_C}$:

$$l \approx \left( \frac{1}{\alpha^{Visual}} - \frac{K}{\alpha} \right)^2 + (K - K^0)^2 \mathrm{P_C} \sqrt{\mathrm{T_C}} + \frac{\mathrm{P_C}}{\sqrt{\mathrm{T_C}}} \left( \frac{1}{\alpha} - \frac{1}{\alpha^0} \right)^2 \quad (9)$$

where $\widehat{w}^{Obs} = \frac{w^{Obs} N^2}{6}$, $\widehat{w}^{\alpha} = \frac{w^{\alpha} N}{2L^2}$, $P_C = \frac{\sqrt{\widehat{w}^{Obs} \widehat{w}^{\alpha}}}{w}$ and $\mathrm{T_C} = \frac{\widehat{w}^{Obs}}{\widehat{w}^{\alpha}}$.

In the 2D case (Fig. 3), $P_C$ and $T_C$ will differ slightly to those derived above, but still describe the same qualitative trade-off between the three terms of the model.

### 5 Neural model of structural inference

#### 5.1 Physiological correlates of message broadcasts and message updates

In the neural implantation of belief propagation, a message broadcast is initiated by the firing of a spike from the broadcasting place cell. This spike is communicated to all neighbouring place cells with conduction delay that is assumed to be inversely proportional to the connection strength (i.e. linearly proportional to the pairwise distance).

In parallel the same spike from the broadcasting place cell $P_B$ drives activity in the grid cell sheet (Fig. 7A) which represents the current belief of the broadcasting place cell's location in grid cell space, since the spike is modulated by the place-to-grid cell weights learned during online exploration. This activity is then assumed to propagate radially outwards at a constant speed from the initial activation site, whilst accumulating noise proportional to the distance travelled.

The development of this activity over time manifests as a travelling wave across the grid cell sheet (Fig. 7B). Mathematically, this wave describes a process of path integration from the initial point emanating radially outward in all directions (since associative relationships between features encode distance not bearing).

The arriving spikes from broadcasting place cells are assumed to produce a sub-threshold depolarisation in receiving place cells ($P_R$), which allows a learning update to its connections to grid cells (representing its belief), shifting connection strength towards currently active grid cells:

$$\boldsymbol{B}_i^{(n)} = \boldsymbol{B}_i^{(n-1)} \cdot \boldsymbol{G}_t \tag{10}$$

i.e. the new belief is simply the product of the current grid cell activity (i.e. the message from the broadcasting place cell; here we refer to a vector of cellular activities, rather than the continuous distribution notation in the main text) and the receiving place cell's prior belief (encoded by the weights $\boldsymbol{B}_i$). Thus, if the spike arrives at the receiving place cell at the same time as the grid cell wave reaches the location to which the place cell was previously associated, there will be relatively little change in the weights (Fig. 7A, left).

Finally, the difference between new and previous beliefs, which is represented by the total change in place-grid cell weights, is calculated in a similar vein to the prediction error term; i.e. using the K-L divergence. This mechanism could plausibly be related to the concentration of a plasticity-related molecule, and the K-L divergence is equivalent to the message tension described earlier in the algorithmic belief propagation solution. Thus, if the learning event produces a significant message tension, the receiving cell will also broadcast its own spike. If the incoming message is not sufficiently different to the current belief, some learning occurs due to the subthreshold input, but no spike is broadcast. This selectivity of firing is necessary to prevent all receiving cells from firing at every time-step, which would produce a travelling front of activity in CA3 as opposed to discrete sequences. Alternatively, receiving a spike could trigger a spike in $P_R$ unless inhibited by coincident activity from the grid cell sheet, as would be the case if the messages were in agreement.

Importantly, the place-place cell weights are not updated during the offline inference process, only during *online* learning. These observations are independent of the current estimate of location such that their encoded pairwise distance estimates decreases in error with experience. Thus, they are the 'ground truth' against which the place-grid cell weights are calibrated.

#### 5.2 Traveling waves in neural media

Traveling waves in neural media have been observed in several brain areas[62] and studied in great detail theoretically[61], often with a focus on their role in synchronizing multiple brain regions. Our model can be viewed as an extension of this theory, describing how plasticity between the HPC and mEC should be affected by precise spike timing. Existing theoretical models of traveling waves and pulses[63,68] assume that the shape of the wave front is unchanged with distance, however our simulations demonstrate that the wave can also be made to broaden with distance, mimicking accumulation of path integration error (Fig. 7B,C).

Traveling waves in mEC are simulated explicitly by calculating the true wrapped distribution at each time-step. Here, we simulate traveling pulses[63], i.e. waves of activity whereby the neural media is only transiently excited (n.b. traveling 'waves' or 'fronts'[61,62] often refer to a radially increasing region of excitation).

Existing traveling pulse solutions preserve the exact shape of the initial stimulus pattern. We developed a model of traveling pulses which broaden with travelled distance, mirroring the accumulation of PI error in all directions. We used a simplified model of wave propagation, which focusses on intuitive demonstrative power rather than biophysical realism (but see Hasselmo, 2014). The proposed model is based on the form of a simple mechanical wave such as oscillations in water:

$$\frac{d\boldsymbol{v}^2}{dt} = c^2\nabla'^2 \cdot H[\boldsymbol{v}] \tag{11}$$

Where $c$ is the speed of wave propagation, $H[\cdot]$ is the Heaviside function and the spatial Laplacian operator:

$$\nabla = \left(\frac{d}{dx}, \frac{d}{dy}\right) = \begin{bmatrix} 0 & 1 & 0 \\ 1 & -4 & 1 \\ 0 & 1 & 0 \end{bmatrix} \tag{12}$$

is replaced by a 2D Gaussian filter with variance equal to the PI noise. It was found empirically that scaling the wave speed $c = \beta c'$ was required to match the desired propagation speed $c$, here $\beta \approx 0.3$.

##### 5.2.1 A note on the effect of adaptation in neural field models

Traveling pulses can also be modelled in with more biophysical detail in neural fields. In a purely excitatory field, propagating activity over time given some measure of agent velocity, while engaging the shifter cells in all directions, would equate to convolving the initial stimulus (e.g. a delta function centred at the grid location of the node from which a message is being sent) repeatedly over time. This would produce a gradually spreading "front" of activity. To produce a traveling "pulse", it is sufficient to introduce to the neurons a linear adaptation (which might represent short-term synaptic depression), which limits the firing of recently active neurons (Ermentrout and McLeod, 1993).

The adaptation rate should be set to the rate of propagation on the sheet. If the propagation and adaptation rates are comparable, a traveling pulse is produced. If the adaptation rate is fast, the region immediately behind the wave will recover quickly.

The latter scenario corresponds to a network implementing path integration via the shifter cell model detailed above; i.e. the mechanism above assumes fast adaptation. Although we do not provide explicit simulations, this qualitative mechanism suggests that the same network might operate in two regimes. At slow propagation speeds (relative to adaptation), the network will perform path integration. At faster propagation speeds, the network may generate traveling pulses using the same circuitry, by setting the head direction signal as uniform across all angles. This qualitative mechanism suggests that "replay", which is implemented in our model by offline traveling waves, necessarily needs to occur at an accelerated rate relative to walking or running.

### 6 Statistical measures

Feature (i.e. place field) locations in the loop closure task were distributed uniformly around the track. For a single sequence $\Theta_s = \{\theta_1^{(s)}, \dots, \theta_{N_s}^{(s)}\}$, the mean sequence length was defined as $\bar{L}_S = \frac{1}{S}\Sigma_{s=1:S} N_s$. The mean hop distance is defined as:

$$\bar{H}_S = \frac{1}{S}\sum_{s=1:S}\sum_{s=2:N_s}\frac{1}{N_s}\mathrm{abs}\left(\left|\theta_i^{(s)} - \theta_{i-1}^{(s)}\right|_c\right) \tag{13}$$

Where $|\cdot|_c$ is the circular distance (minimum of the clockwise and anticlockwise distances).

The 'structural error' is defined as the mean absolute difference between the true pairwise distances and those pairwise distances recovered through *offline* inference:

$$\epsilon_{pairwise} = \frac{1}{|E|}\sum_{(i,j)\in E}\left|\delta_{ij} - d\left(\boldsymbol{b}_i^{max}, \boldsymbol{b}_j^{max}\right)\right| \tag{14}$$

where $\boldsymbol{b}_i^{max}$ is either the unique feature encoding on the grid sheet (*metric* space) resulting from maximum-likelihood solutions to the *offline* model, or the peak of the landmark distribution recovered during belief propagation.

### 7 Analysis of shearing due to inhomogeneous environmental input

#### 7.1 Structure of the PC→GC weights

In simulation, the PC→GC weights are learned via the BCM rule (Online Methods, Eq. 9). The steady-state weights in the case where the firing of both cells is independent approximates the product of their spatial firing patterns. In 1D, this corresponds approximately to the integral (assuming an infinitely long track):

$$B_{pg} = \int_{-\infty}^{\infty} P_p(\bar{x}) G_g(\bar{x}) d\bar{x} \tag{15}$$

Where the constant of proportionality is the $p^{th}$ diagonal entry of the place field's precision matrix. We assume the firing pattern of grid cell $G_g(\bar{x})$ over physical space $\bar{x}$ is approximated as the sum of multiple Gaussians with mean separated periodically across space:

$$G_g(\bar{x}) = \sum_{i=-\infty}^{\infty} N(\bar{x}, \mu_{ig}, \sigma_g^2) \tag{16}$$

Where $\mu_{ig} = i\lambda_g$ and $\sigma_g^2$ are the centre of mass and tuning width of the grid fields respectively and $\lambda$ is the grid scale. Place cells are also modelled as Gaussians:

$$P_p(\bar{x}) = N(\bar{x}, \mu_p, \sigma_p^2) \tag{17}$$

Where $\mu_p$ is the preferred centre of firing of a given place cell in real space. The product of two Gaussians is also Gaussian such that:

$$B_{pg} = \int_{-\infty}^{\infty} \sum_{i=-\infty}^{\infty} S_{pg_i} N_{pg_i}(\bar{x}, \mu_p + \mu_{g_i}, \sigma_p^2 + \sigma_g^2)\, d\bar{x} = \sum_{i=-\infty}^{\infty} S_{pg_i} \tag{18}$$

Where:

$$S_{pg_i} = \frac{1}{\sqrt{2\pi(\sigma_p^2 + \sigma_g^2)}} exp\left[-\frac{(\mu_p - \mu_{g_i})}{2(\sigma_p^2 + \sigma_g^2)}\right] \tag{19}$$

#### 7.2 Reweighting the place cell activity

Given the steady-state PC-GC weights learned above, the *observation* model output for a given grid cell *g* would be given by the weighted sum of the place cell activity across space. To model a bias in saliency (Section 2, Fig. S3), we next introduce a reweighting function that corresponds to a linear but spatially non-uniform scaling of the firing rates (or equivalently, synaptic weights). The re-weighting function $w(\mu_p)$ was chosen arbitrarily to have simple exponential-quadratic form, to capture simple patterns of inhomogeneity in place cell input:

$$w(\mu_p) = e^{r_0 + r_1(\mu_p + \bar{x}_c) + r_2(\mu_p - \bar{x}_c)^2} \tag{20}$$

Where the grid pattern is being shifted relative to some focal point of shifting $\bar{x}_c$ according to some quadratic coefficients $\{r_0, r_1, r_2\}$.

Assuming the number of place cells is large and uniformly distributed and the receptive field widths of the place and grid cells are equal $\sigma_P^2 = \sigma_g^2 = \sigma^2$, we analyze (for simplicity) the effect of the reweighting on a single grid field, assuming that the interaction between neighbouring grid fields is negligible. The un-normalized output from the observation model is:

$$H_g(\bar{x}) = \int_{-\infty}^{\infty} p(\mu_p, \bar{x}) \cdot B_g(\mu_p, \sigma^2) \cdot R(\mu_p) d\mu_p \tag{21}$$

During *online* localization (Main Text Eq. 2), the total grid cell activity at each time-step is normalized. Thus, we can solve for the peaks of the re-weighted and normalized observation model output model output $\hat{H}(\bar{x}, \mu_g) = H(\bar{x}, \mu_g) / \int H(\bar{x}, \mu_g) d\mu_g$ (again assuming large number of grid cells) by setting $\frac{dH(\bar{x}, \mu_{g_i})}{dx} = 0$ results in:

$$\delta\bar{x} = \sigma^2 \left(-r_1^2 + 2r_2(\bar{x}_c - \mu_g)\right) \tag{22}$$

Thus, applying the re-weighting produces a shift in the grid fields $\mu_g$ which is linearly proportional to the distance from the grid field to the centre of shearing $\bar{x}_c$, and scaled by the magnitude of the reweighting parameters and the tuning width $\sigma^2$. $r_1$ produces a constant offset, and so does not affect the magnitude of shearing. We therefor simplify the above to:

$$\delta\bar{x} = \sigma^2 R\left(\bar{x}_c - \mu_g\right) \quad (23)$$

Where $R = r_2$ corresponds to re-weighting strength in Main Text Fig. S3B. A similar analysis can be performed in 2D, with a reweighting function $w(\boldsymbol{\mu}_p) = e^{R(\mu_{px}-\bar{x}_c)(\mu_{py}-\bar{y}_c)}$ to obtain the relationships:

$$\delta\bar{x} = \sigma^2 R(\bar{x}_c - \mu_{gy}) \quad (24a)$$
$$\delta\bar{y} = \sigma^2 R(\bar{y}_c - \mu_{gx}) \quad (24b)$$
